# mEMbrain: an interactive deep learning MATLAB tool for connectomic segmentation on commodity desktops

**DOI:** 10.1101/2023.04.17.537196

**Authors:** Elisa C. Pavarino, Emma Yang, Nagaraju Dhanyasi, Mona Wang, Flavie Bidel, Xiaotang Lu, Fuming Yang, Core Francisco Park, Mukesh Bangalore Renuka, Brandon Drescher, Aravinthan D.T. Samuel, Binyamin Hochner, Paul S. Katz, Mei Zhen, Jeff W. Lichtman, Yaron Meirovitch

## Abstract

Connectomics is fundamental in propelling our understanding of the nervous system’s organization, unearthing cells and wiring diagrams reconstructed from volume electron microscopy (EM) datasets. Such reconstructions, on the one hand, have benefited from ever more precise automatic segmentation methods, which leverage sophisticated deep learning architectures and advanced machine learning algorithms. On the other hand, the field of neuroscience at large, and of image processing in particular, has manifested a need for user-friendly and open source tools which enable the community to carry out advanced analyses. In line with this second vein, here we propose mEMbrain, an interactive MATLAB-based software which wraps algorithms and functions that enable labeling and segmentation of electron microscopy datasets in a user-friendly user interface compatible with Linux and Windows. Through its integration as an API to the volume annotation and segmentation tool VAST, mEMbrain encompasses functions for ground truth generation, image preprocessing, training of deep neural networks, and on-the-fly predictions for proofreading and evaluation. The final goals of our tool are to expedite manual labeling efforts and to harness MATLAB users with an array of semi-automatic approaches for instance segmentation. We tested our tool on a variety of datasets that span different species at various scales, regions of the nervous system and developmental stages. To further expedite research in connectomics, we provide an EM resource of ground truth annotation from 4 different animals and 5 datasets, amounting to around 180 hours of expert annotations, yielding more than 1.2 GB of annotated EM images. In addition, we provide a set of 4 pre-trained networks for said datasets. All tools are available from https://lichtman.rc.fas.harvard.edu/mEMbrain/. With our software, our hope is to provide a solution for lab-based neural reconstructions which does not require coding by the user, thus paving the way to affordable connectomics.

## 1. Introduction

Connectomics, the spearhead of modern neuroanatomy, has expanded our understanding of the nervous system’s organization. It was through careful observation of the neural tissue that Santiago Ramon y Cajal, father of modern neuroscience and predecessor of connectomics, reasoned that the nervous system is composed of discrete elements - the nerve cells. He further hypothesized key functional cell and circuit properties, such as neuronal polarity and information flow in neuronal networks, from anatomical observations, documented in extraordinary drawings. Connectomics - in particular based on electron microscopy images - has progressed immensely, and while the first complete connectome - the “mind of a worm” - was a manual decade-long endeavor for a reconstruction of merely 300 neurons (1), technological and methodological strides have enabled the field to elucidate complete circuitry from several other neural systems (2–12).

It would be highly impoverishing to view connectomics’ purpose as merely the pursuit of neural circuit cataloguing. In recent years, in fact, connectomic reconstructions have been a new tool instrumental to answering outstanding questions in various subfields of neuroscience, which required synaptic resolution. Developmental studies have vastly benefited from microconnectomic reconstructions, opening the possibility of investigating precise synaptic rearrangements that take place in the first stages of life (11, 13–15). Further, circuit reconstructions have allowed in-depth studies of phylogenetically diverse systems, such as the ciliomotor system of larval *Platynereis*, (16, 17), learning and memory in *octopus vulgaris* (18), the olfactory and learning systems of *Drosophila* (10, 19) and the visuomotor system of *Ciona* (20). Connectomes have also provided insights into systems neuroscience, where avenues to pair structural and functional data from the same region of the brain are being explored. Noticeable examples of such endeavors are the study of mechanosensation in the zebrafish (21), the study of the posterior parietal mouse cortex, important for decision making tasks (22), and the functional and structural reconstructions of a mouse’s primary visual cortex (23–25). Further, connectomes have proven to be a useful - and perhaps necessary - resource for computational modeling and simulation of circuits, by providing biological constraints such as connectivity, cell types and their anatomy. For example, the fly hemibrain (10) was queried to find cell candidates performing specific neural computations (26), murine connectomes have been shown to allow for discrimination between different candidate computational models of local circuits (27), and the *C. elegans* connectome is being leveraged to simulate the first digital form of life through the open science project “OpenWorm” (28). Finally, we are at an exciting moment in connectomics’ history, as recent reconstructions allow us to open a window on the human brain (29). This important milestone, in conjunction with contemporary efforts to develop a whole mouse connectome (30), will enable the community to reconstruct circuits in the context of neuropathology, and shed light on wiring diagram alterations that give rise to the so-called “connectopathies” (30–32).

All these neural reconstructions have become a reality due to the progress in tissue preparation for electron microscopy and the tremendous progress in computer vision and artificial intelligence techniques. On the one hand, progress in tissue staining, cutting, imaging and alignment has yielded traceable volumes amenable to reconstruction. However, manual reconstructions alone of neural circuits represent a massive endeavor. Previous studies have computed that manual reconstruction of medium-sized neural circuits would amount to hundreds of thousands of hours of human manual labour, and would be a multi-million dollar investment, which is a prohibitive effort in most settings (33). Further, this manual effort quantification is highly variable depending on the precision requested by the research question at hand - highly precise annotations require a quasi-pixel accuracy, which naturally lengthen the time of the procedure. Therefore, manual annotation alone is not scalable for entire neural circuit reconstruction. Recently, machine and deep learning techniques have become of common use for segmenting neural processes, thus aiding and expediting hefty manual annotation, and paving the way to high-throughput neural architecture studies. In this frame, convolutional neural networks (CNNs) have emerged as a successful solution for pixel classification. A typical automatic neurite reconstruction first begins by inferring probability maps of each pixel/voxel in the image for classifying boundaries of distinct cells (34, 35). In particular, U-net architectures have become common practice for biomedical image segmentation (36), and are widely employed to achieve this first task. In a second step, a different algorithm agglomerates the pixels/voxels confined within the same cell outlines.

In the recent years, the field has benefited from deep learning algorithms designed specifically for the task of connectomic instance segmentation on particularly large and challenging datasets. One notable example of this is the Flood Filling Network architecture, a 3D CNN paired with a recurrent loop which segments in the volume one cell at a time by iteratively predicting and extending the cell’s shape (37). A similar end-to-end approach iteratively segmenting one cross section of a neuron at a time has been pursued independently (38). Recently this approach has been extended by training networks to flood fill numerous objects in parallel (39). Many of these elaborate and heavily engineered pipelines (see also the Supplementary Material Section 1.2) present open source code repositories, however they remain of difficult practical use for researchers who do not have a software or computational background. For these reasons, many of the largest connectomics efforts have been carried out in collaboration with teams of computer scientists or even companies, option that requires a great deal of resources, both in terms of funding, and in terms of computing and storage capabilities.

While on one hand it is imperative to ever better the accuracy and scalability of these advanced algorithms, the field of image processing in particular, and science at large, have felt the urge for more democratic and easily accessible tools that can be intuitively employed by independent scientists. To name a few, tools such as ImageJ for general and multi-purpose image processing (40, 41), Ilastik (42) and Cellpose (43) for cell segmentation, suite2p for calcium imaging (44), Kilosort for electrophysiological data (45), DeepLabCut (46) and Moseq (47) for behavioral analyses have enabled and empowered a larger number of scientists with the ability to carry out significant studies that previously would have been challenging or unfeasible, requiring non-trivial technical skills, time and resources. More specifically to the field of connectomics, there are a plethora of open software, mostly geared towards image labeling for manual reconstruction. Examples include but are not limited to VAST lite (48), Ilastik (42), NeuTU (49), Knossos (50) with its online extension webKnossos (51), and Reconstruct (52). Because most of these software tools do not include a deep learning-based segmentation pipeline, a few software packages have been proposed to supply a CNN-based reconstruction, such as SegEM (33) which relies on skeletonized inputs for example from Knossos, and Uni-EM, a python-based software that wraps many of connectomics’ image processing techniques (53).

We reckoned that making connectomics an affordable tool used by single labs meant providing a desktops solution compatible with the most common operating systems and computational frameworks currently used in the field. Thus, we focused our efforts here on creating a package based on MATLAB, which is one of the most commonly used coding environments in the basic science communities, providing its users with a rich array of image processing and statistical analysis functions. Importantly, our main task here was not to present new functions for computer vision for connectomics, but rather we propose existing functions and machine learning models in a simple and user-friendly software package. Hence, the accuracy of our tool derives from the solutions presented previously in connectomics. As a second goal for our tool, we wished to create a virtuous and rapid EM reconstruction cycle which did not require solving the more expensive automated reconstruction problem. Thus, our deep learning tool greatly accelerates manual reconstruction in a manual reconstruction framework called VAST (48), an annotation and segmentation tool widespread in the community with numerous tools and benefits for data handling and data visualization. We expect our ML tools to be valuable to researchers that already use VAST. Thus, we created mEMbrain, a segmenting tool for affordable connectomics with the following attributes:

- mEMbrain has an interactive, intuitive, and simple interface, which leverages image processing and deep learning algorithms requiring little to no coding knowledge by the user.
- mEMbrain is a MATLAB-based extension of VAST, a segmentation and annotation tool widely used in the Connectomics community (48). Using VAST as a server proves to be a clear-cut solution as it can splice the data and cache the space on demand, allowing mEMbrain to run on any cubical portion of datasets, independently of how the images are stored at the back-end.
- mEMbrain processes datasets locally on commodity hardware, thereby abolishing the need of expensive clusters and time-consuming data transfers.

We validated the robustness of mEMbrain by testing it on several species across different scales and parts of the nervous system in diverse developmental stages, and demonstrated mEMbrain’s usefulness on datasets in the terabyte range. Further, we tested mEMbrain’s speedup in terms of manual annotation time, and observed several fold improvement in manual time. All together, this paper presents new connectomic tools in platforms that had poor support for connectomic research. Furthermore, our tool extends the functionality of VAST to allow semi-automated reconstruction, already offered by other platforms.

## 2. mEMbrain’s Concept

mEmbrain is a software tool that offers a pipeline for semi-automatic and machine learning-aided manual reconstruction of neural circuits through deep convolutional neural network (CNN) segmentation. Its user interface guides the user through all the necessary steps for semi-automatic reconstruction of electron microscopy (EM) datasets, comprising ground truth generation with data augmentation, data preprocessing, CNN training and monitoring, predictions based on electron microscopy datasets loaded in VAST, and on-the-fly validation of such predictions in VAST itself. mEMbrain is written in MATLAB, in order to interface seamlessly with VAST, a widely used annotation and segmentation tool (48). Most of today’s pipelines involving machine and deep learning rely on Python, which although incredibly proficient and widely used in the computational community, is still less adopted in biological fields. We wanted to bridge this gap to make connectomics more accessible to a larger biological science community. mEMbrain can run on any operating system where both VAST and MATLAB (with parallel computing and deep learning toolboxes installed) are operative.

mEMbrain is a democratizer of computational image processing, which is necessary for EM circuit reconstruction. Its main purpose is to collect functions and processes normally carried out by software or computational scientists, and to embody them in a single software tool, which is intuitive and user-friendly, and accessible to any scientist. Thus, no coding skills are required for mEMbrain’s operation.

mEMbrain’s practicality starts from its installation. In many cases, software installation represents a hurdle, which in turn makes the frustrated user disinterested. To ease installation, our tool is a 352 KB folder downloadable from our GitHub page (github/mEMbrain). Once running, mEMbrain hosts all of its tools in one unique interface designed to be intuitive and user-friendly. mEMbrain’s design is modular, with every tab presenting a different step of the workflow. Thus, the user can either be guided through the pipeline by following the tab order, or they can access directly the processing step of interest. For more information on mEMbrain’s download and setup, please see the Supplementary Materials section 1.1.

The main concept of mEMbrain is to create a synergistic dialogue with VAST in order to automate parts of the connectomic pipeline (see Figure 1). Typically, VAST is adopted by researchers for electron microscopy annotation and labeling. Such labeling and labeled microscopy can then be exported and directly used in mEMbrain, an independent package that complements VAST and VastTools by applying machine learning algorithms on image data. Within mEMbrain, images are processed and used to create datasets for training a deep learning model for semantic segmentation. Other than evaluating the results of the training phase through learning curves, the researcher can directly test how well the trained model performs, by making predictions on (portions of) the EM dataset open in VAST. The predictions are visible on-the-fly in VAST, superimposed on the open dataset. If the results achieved are not satisfactory, the user can improve the model by providing more ground truth examples; it is especially beneficial if the new labels incorporate regions and features of the dataset where the model predicted poorly. Hence, the newly generated ground truth is incorporated in the training dataset, and the deep learning model is retrained. This iterative process is continued until the results are deemed appropriate for the task at hand. In some cases, the iterative generation of new ground truth can be accelerated by making the deep learning segmentation editable in VAST, so that the researcher can swiftly correct such segmentation, saving time. Finally, once the prediction result is satisfactory, the final semantic segmentation can be leveraged for accelerating neural circuit reconstruction, by either using the predictions as a VAST layer, which dramatically speeds up manual painting (the main use-case of mEMbrain), using border predictions or by performing 2D instance segmentation. 3D instance segmentation algorithms are currently not incorporated in mEM-brain, but can be used in synergy with mEMbrain as surveyed in Section 4 and the supplementary material (Section 1.2 and Figure S1). This utility to encompass all the machine-learning steps within mEMbrain as part of 3D instance segmentation algorithms was recently demonstrated in a study of human brain biopsies for connectomics (see (32) and reconstruction at https://lichtman.rc.fas.harvard.edu/mouse_cortex_at_1mm).

**Fig. 1.**
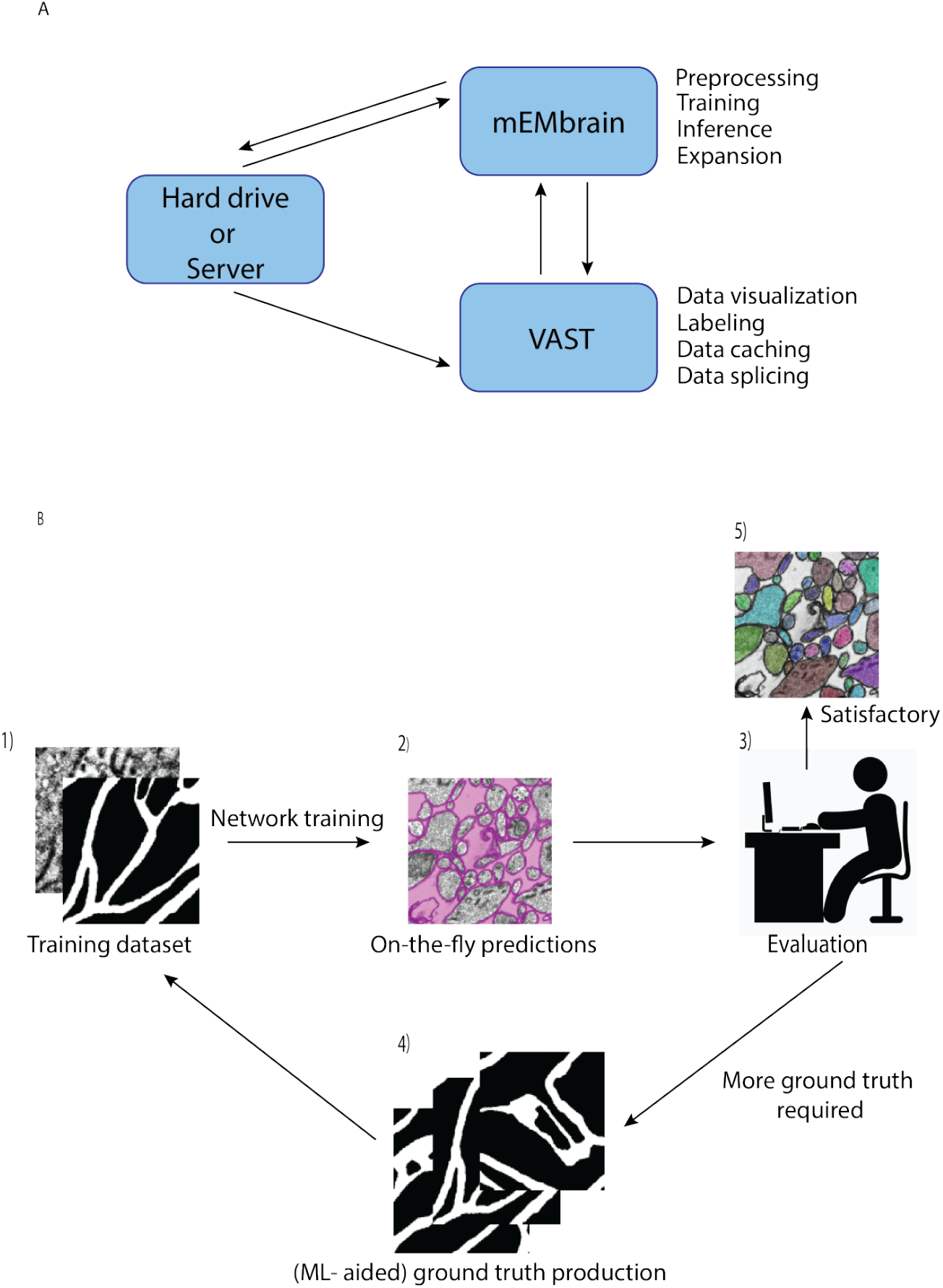
mMEbrain’s workflow and integration with VAST. **(A):** communication between mEMbrain, VAST and data storage. mEMbrain and VAST communicate bidirectionally, as VAST stores, caches and splices the data which can then be imported into mEMbrain through VAST’s application programming interface. mEMbrain’s outputs are then transferred back to VAST for visualization and postprocessing. mEMbrain can also access the data directly at where it is stored, and will save there its outputs (if so the user indicates). **(B):** mEMbrain’s iterative workflow. The user starts by creating a training dataset of EM and corresponding labels 1), which are then used to train a convolutional deep learning network. The results of such network can be visualized on-the-fly directly on datasets open in VAST 2). Further, if the researcher is satisfied with the current state of network inference, they may proceed to a semi-automatic approach for semantic segmentation 3). However, if they are not satisfied with the current output of the network, the can use these predictions to accelerate further ground truth production 3-4), which is then incorporated in further training of the network to achieve better results 4).

As a technical note on mEMbrain’s hardware requrirements, there are none beyond what is required for the installation of MATLAB and VAST. The memory footprint of mEMbrain does not exceed the amount needed for the operation of MATLAB and VAST alongside the memory requirements to iteratively read small chunks of the image space. mEMbrain writes to disk the predicted images as 1024×1024 PNGs without buffering, in a format consistent with VAST’s tiling of the image space. If electron microscopy images are also chunked in VAST into 1024×1024 pixel images, then for each of the input images and output channels this buffer will be the overhead RAM requirement for mEMbrain (i.e., an order of MBs).

## 3. Example Workflow

We here report the various steps of the image processing pipeline we have implemented and wrapped within mEMbrain. For the typical flow, refer to Figure 2 for our general purpose GUI, and Figure 4 for the specific pipeline adopted for the *C. elegans* data described in Section C.

**Fig. 2.**
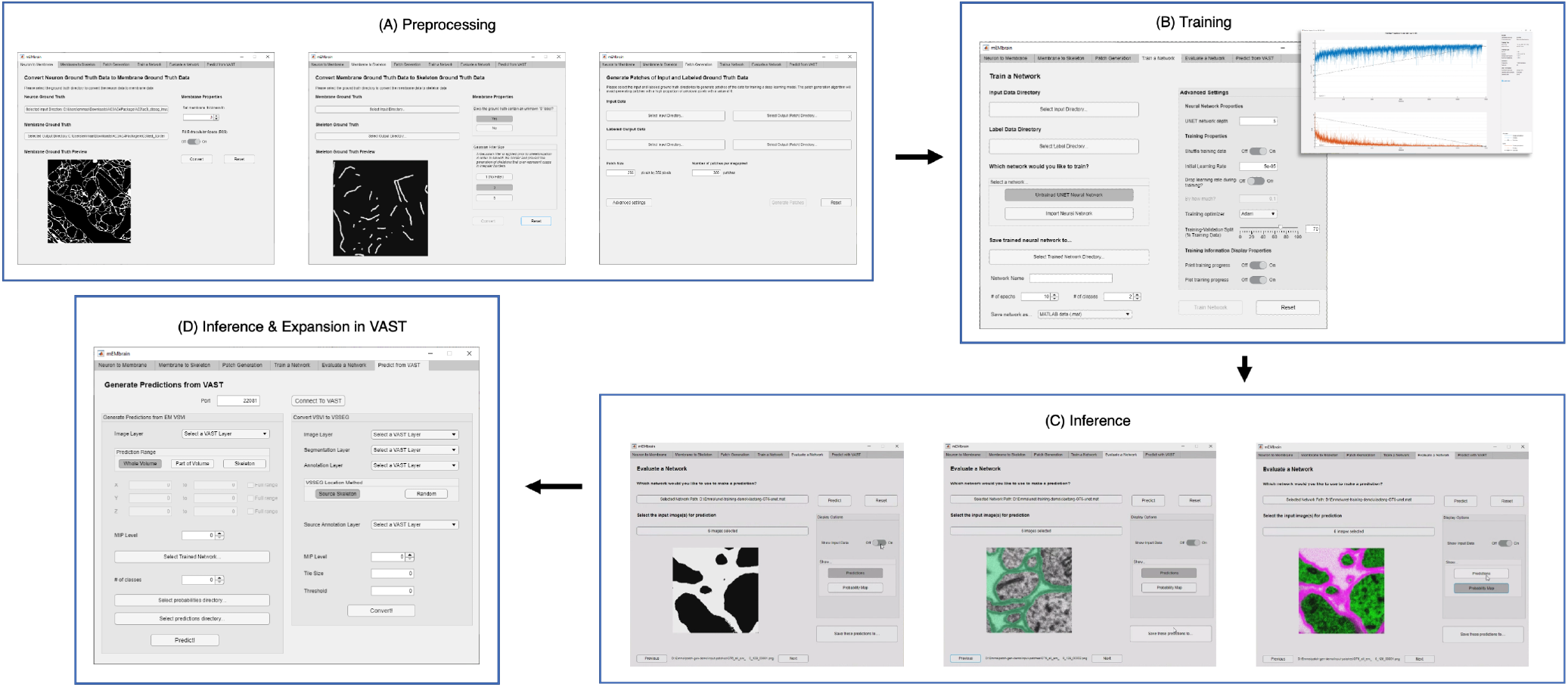
mEMbrain’s MATLAB-based GUI for data preprocessing, training, inference, and integration with VAST. All the functions are collected in one user interface, and can be accessed by clicking the different tabs. **(A)**: These first 3 tabs allow the user to create a training dataset from EM images and corresponding ground truth. **(B)**: Further, the user can train a deep neural network. As default, we make use of MATLAB’s built-in U-net, whose training can be customized through the various user-chosen parameters. The training’s progress can be monitored by MATLAB’s learning curves. **(C)**: To evaluate a network, predictions can be made on small sample images, as seen in the small squares of the GUI. **(D)**: Finally, researchers can infer directly on-the-fly in VAST on the dataset herein open. Further, they can convert such inference to editable layers in VAST, that may be leveraged for machine learning (ML)-based ground truth preparation.

### A. Dataset creation and image preprocessing

The first step towards training neural networks for segmentation is the creation of a training dataset composed of both images and associated ground truth, or labels. It is common wisdom that abundant ground truth will yield a better prediction of the training algorithm. We realize that the preparation and curation of a comprehensive training dataset can represent a hurdle for many researchers. One strategy might be to label many EM images; however, this requires many hours dedicated to tedious manual annotation. Alternatively, computational methods can be leveraged for augmenting ground truth with image processing techniques - hence necessitating less labeling; however, this requires having a good mastery of coding skills. Thus, we incorporated a dataset creation step, which allows researchers to process the labeled images paired with their EM counterpart with just a few mouse clicks. Once the user has imported the microscopy images coupled with their labels, mEMbrain converts the latter in images with 2 or 3 classes, depending on the task at hand. The EM images are then corrected by stretching their grayscale. Subsequently, patches of a user-chosen dimension are extracted from the pair of EM and label images. Notably, mEMbrain first verifies the portion of the image that presents a saturated annotation (i.e. areas where every contiguous pixel is annotated), which can assume any arbitrary shape desired by the researcher. Then, mEMbrain efficiently extracts patches from such regions. Thus, the images do not have to be fully annotated for them to be incorporated in the training dataset, and this feature makes the region of interest selection more flexible, faster and seamless.

One noteworthy feature of this step is the incorporation of data augmentation, in the hope that fewer annotated images are required to obtain a satisfying result. In particular, we verified that rotations yielded a better result during testing phase, hence we implemented a random rotation of any possible degree for every pair of patches. At each rotation of the ground truth data, mEMbrain uses the chessboard distance (or Chebyshev distance) between labeled pixels of the ground truth to the closest unlabeled pixels. Then, mEMbrain individuates pixels around which a square patch of user-defined size will contain fully annotated pixels, and such pixels are then used for patch generation. Further data augmentation methods such as image flipping, Gaussian blurring, motion blurring and histogram equalizer are also implemented. This ensemble of techniques ensures that nearby regions from the same image can be more heavily sampled for patch generation without making the training overfit such a region, allowing the extraction of “more patches for your brush stroke”. It is important to note that this feature is one of the only algorithmic novelties of mEMbrain. Hence, although we are not in the position to benchmark the networks’ performances (as the U-net architecture is not our contribution), we here show that our patch generation and data augmentation is as good - if not better - as many off-the-shelf methods (see Figure 3).

**Fig. 3.**
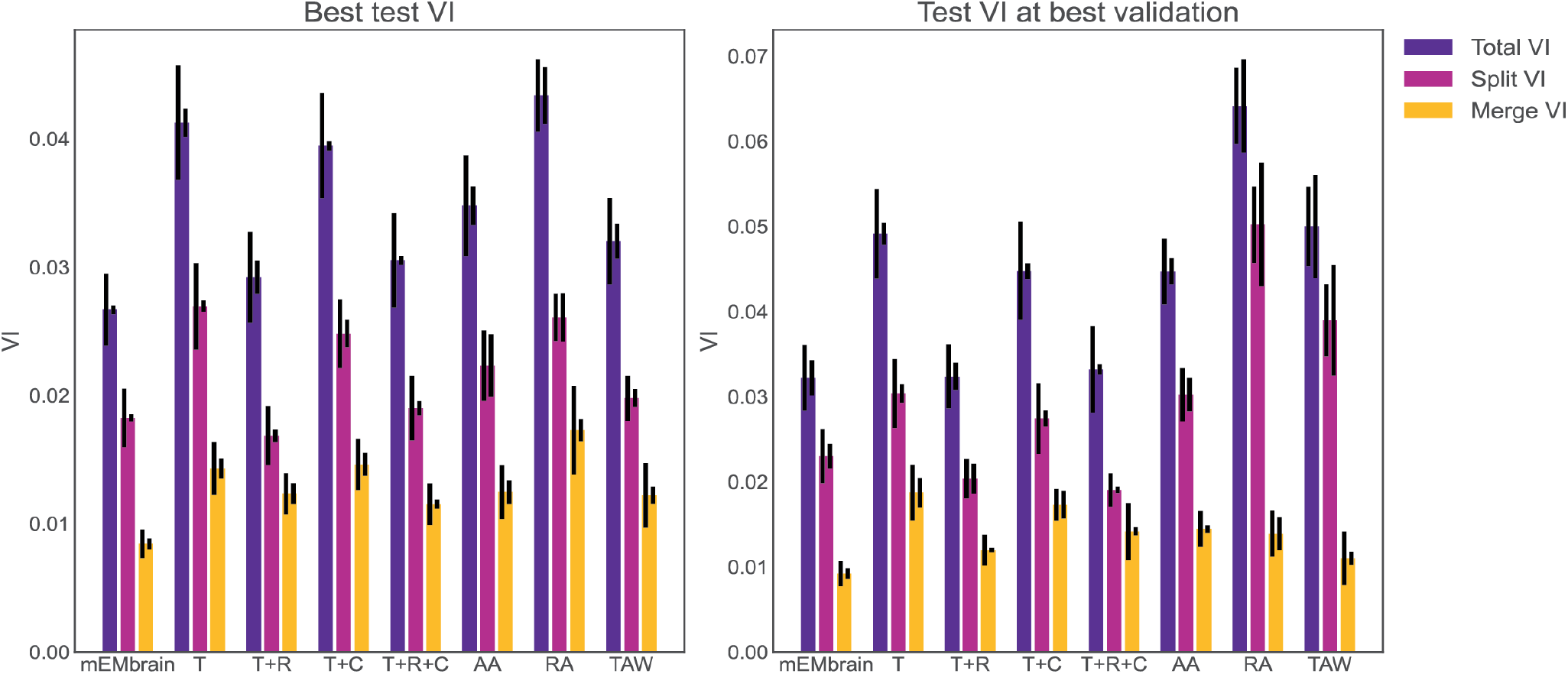
Evaluation of mEMbrain’s data augmentation. (Left) The best achieved test variation of information (test VI) for different augmentation methods. The left error bar is the standard error of mean for 6 different test images. The right error bar is the standard error of mean when re-training the network with different seeds. We observe that mEMbrain achieves a better(lower) test VI when compared to simple augmentation methods and published general methods. (Right) Same as left but we plot the test variation of information for different augmentation methods for the network producing the best validation loss. The abbreviations stands for T: translation only. T+R: translation and rotation. T+C: translation and color jitter augmentation. T+R+C: translation, rotation and color jitter augmentation. AA: AutoAugment (54). RA: RandAugment (55). TAW: Trivial Augment Wide (56).

**Fig. 4.**
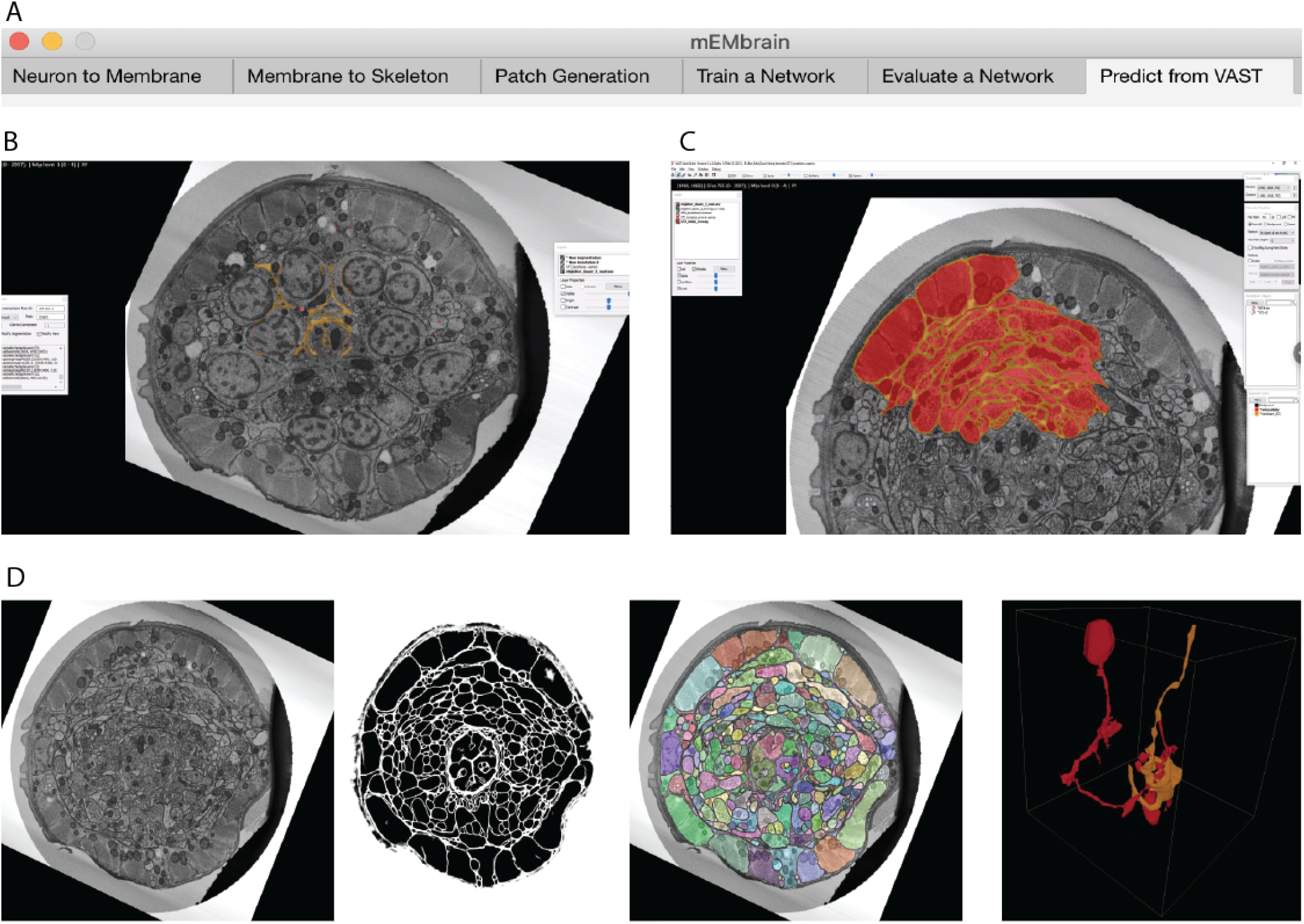
Example workflow with mEMbrain and VAST. **(A)**: mEMbrain’s header with step-by-step pipeline for deep learning segmentation of EM images. **(B)**: The user can initially apply a pretrained network on the dataset at hand and use these predictions both for a) a first evaluation of which areas of the dataset should be included in ground truth, and b) as a base for ground truth generation. The figure shows VAST’s window with the *C. elegans* dataset open. In orange, the predictions of a pretrained network are shown. **(C)**: The user can convert the predictions to an editable layer in VAST, and use these as rough drafts of ground truth. By manually correcting these, one can generate labels in a swifter manner, saving a significant amount of time (roughly half in the case of the *C. elegans* dataset). **(D)**: Left: EM section of the *C. elegans* dauer state dataset. Second panel: mEMbrian’s predictions of the same section. Third panel: mEMbrain’s 2D expansion. Right: example of 3D reconstruction obtained through automatic agglomeration algorithms. The two reconstructed neurites are shown in VAST’s 3D viewer.

**Fig. 5.**
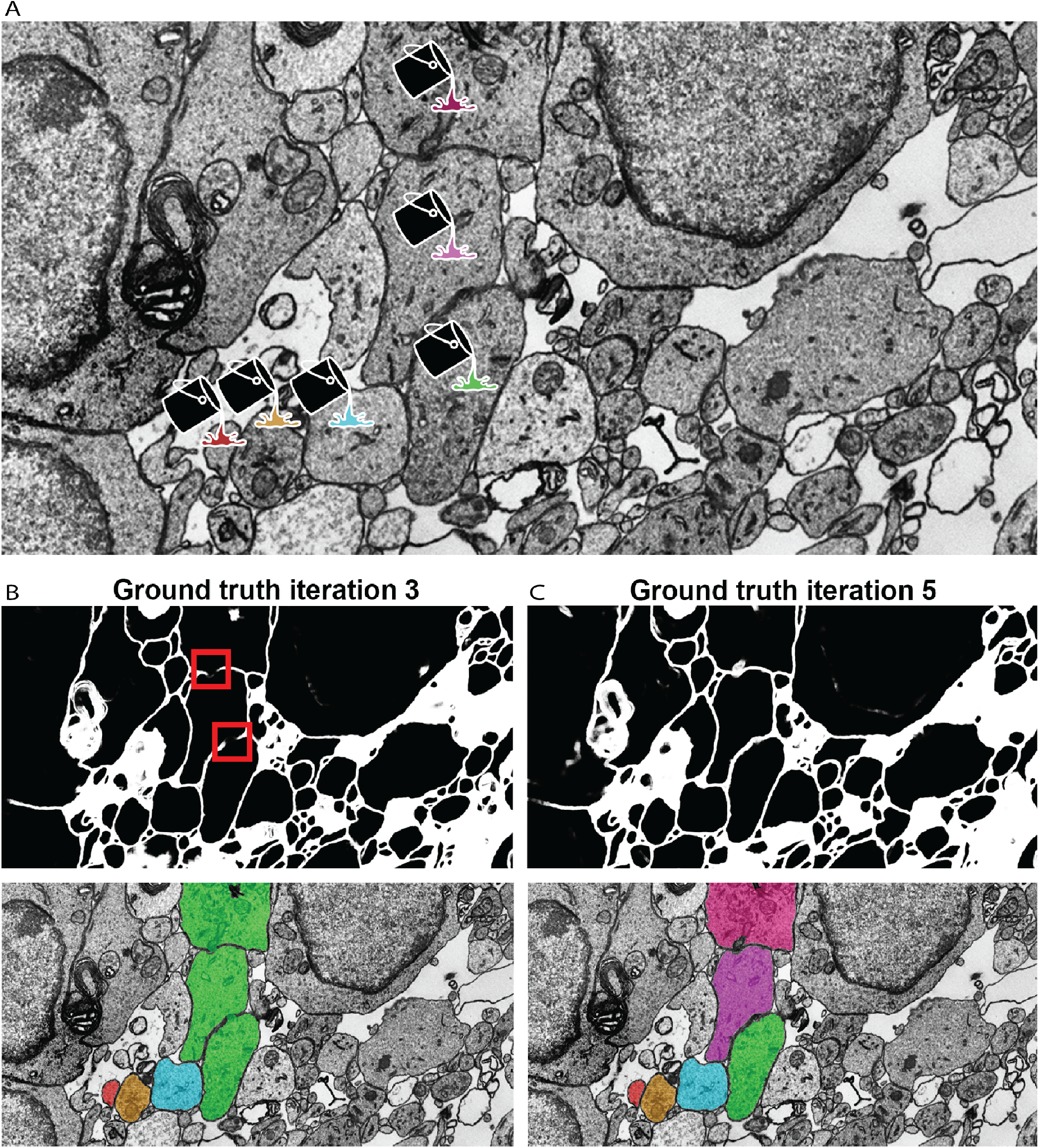
Network qualitative assessment via membrane predictions. **(A)**: EM image opened in VAST. When using the membranes to constrain VAST’s flood filling functionality (see Section D), the virtual paint will fill every pixel which is contained within the constraining boundaries (here, the membranes). Thus, if the predicted membranes are broken as in panel **(B)**, these will lead to so-called merge errors, where multiple different cells are labeled with a same color and ID. Broken membranes are a symptom of a poorly trained network, and hence it may benefit from more training with further ground truth, as seen in panel **(C)**.

In addition to this “smart” patch generation feature, mEMbrain also includes conversion features for a) instance segmentation ground truth to contours ground truth (i.e. membrane ground truth) and b) membrane ground truth to skeleton ground truth. The former uses erosion and dilation with a user-specified filter radius to transform filled-in neuron segmentation annotations into membrane ground truth of a specified thickness. mEMbrain can also generate this membrane data with or without extracellular space filled in. For the latter feature, mEMbrain uses MATLAB’s built-in 2D binary skeletonization functions to generate 2D neuronal skeletons from membrane ground truth. The utility of such ground truth conversion lies in the possibility to then train subsequent deep learning networks in a supervised manner to learn and predict the medial axis of the neuronal backbone. Learning such neuronal backbone enhances the ability of existing reconstruction algorithms to agglomerate objects, as seen in the supplementary material (Section 1.2). While we implemented agglomeration techniques using neuronal backbones predicted by EM (see Figure S1), these will be integrated into mEMbrain’s subsequent software release, and are here described for their novelty and to allow the connectomics community to further test and explore such methods.

### B. Network training

Once datasets are created and preprocessed, researchers are in the position to train a network for image segmentation. There are two options for approaching the training phase:

1. train a pre-implemented U-net (36);
2. load a pre-defined network and continue training upon it.

The implementation of U-net was chosen given the success this deep learning architecture has in the field of biomedical imaging segmentation. Although the implementation of the network is built-in to MATLAB, the user still preserves ample degrees of freedom for customizing parameters of both the network architecture - such as the number of layers - and the training - for example the hyper parameters and the learning algorithm. As of now, most of mEMbrain’s features work when predicting 2 or 3 classes for the step of semantic segmentation. Importantly, the network can be saved as a matrix with trained weights, which can then be used for future transfer learning experiments (see section F).

Alternatively, pre-trained networks can be loaded in mEMbrain to be re-trained. As discussed in Section F, learning upon a pre-trained network, a strategy in the domain of transfer learning here referred to as *continuous learning*, typically yields better results with less ground truth. Of note, it is possible to import networks that have been trained with other platforms, such as PyTorch or Tensorflow, thanks to designated MATLAB functions (for a tutorial, the reader is referred to (57)), or to import/export trained networks and architectures using the ONNX (Open Neural Network Exchange) open-source AI ecosystem format which is supported by various platforms including mEMbrain and MATLAB. Since much of deep-learning-enabled connectomics is done in Python-based machine learning platforms, we also wanted these users to be able to integrate mEMbrain into their workflows. Thus, we also implemented a feature where users can export neural networks trained on ground truth data in mEMbrain to the Open Neural Network Exchange (ONNX) format. This format preserves the architecture and trained weights of the model, allowing the user to import the model back into Python-based platforms such as Tensorflow and Pytorch for further investigation and analysis. In terms of training time, training of 6702 patches took 32 minutes, which amounts to an average of 13.95 patches/second on a Nvidia RTX 2080Ti GPU.

For a quick assessment of deep learning model training, we implemented an evaluation tab, where one can use such a model to make predictions on a few test images and qualitatively gauge the goodness of the network.

### C. On-the-fly predictions with VAST

Once one has trained a deep learning model for semantic segmentation and is satisfied with its results, prediction on the dataset may be carried out. mEMbrain has 3 different modalities for prediction, namely:

- predictions on whole EM volumes;
- predictions on specific regions of the EM volume;
- predictions around anchor points positioned in VAST.

When users predict on whole EM datasets - or portions of it - by either inserting the coordinates delimiting the regions of interest or by using VAST’s current view range, mEMbrain requests EM matrices from an image layer in VAST through the application program interface (API). Our implementation speeds up EM exporting by optimizing the image request and tailoring it to VAST’s caching system (48). Because VAST caches 16 contiguous sections at one given time, mEMbrain requests chunks of 16 [1024 × 1024] sections at a time, reading first in the dataset’s *z* dimension, proceeding then in the *x* and *y* dimensions. Data is read at the mip level chosen by the user, which should match the resolution at which the network was trained. Once having read the EM images, mEMbrain corrects them with the same grayscale correction that was applied when preparing training datasets, and then it predicts the semantic segmentation with the chosen deep learning model. Because the training phase occurs on patches that have dimensions in multiples of [128 × 128] pixels, predictions on [1024 × 1024] pixels at a time is a valid operation. Once image pixels are classified, the predictions are saved as .pngs in a folder designated by the user. At the same time, mEMbrain creates a descriptor file (with extension .vsvi), which is a text file following the JSON syntax that specifies the naming scheme and the storage location of the predicted images, as well as other metadata necessary for the dataset. Once created, the .vsvi file can be loaded (dragged and dropped) in VAST, which then loads the predictions, which can be viewed superimposed on the EM dataset. For best VAST performances and smooth interactions with MATLAB, we recommend having a RAM of at least 64 GB.

It might be useful, in some scenarios, to predict and segment only particular regions which do not all align along the same *z* axis. Leveraging VAST’s skeleton feature, researchers may allocate anchor points in regions of interest throughout the dataset. mEMbrain can then predict locally around such anchor points. One example of such scenario is when trying to determine if a deep learning model provides satisfactory predictions on a large dataset. For such evaluation, mEMbrain can predict a set of cubes centered around pre-selected coordinates (represented by VAST skeletons). Based on the outcome, the user can decide if the model’s output is satisfactory. Another example is the prediction only around certain regions of interest sparse through the dataset, such as synapses.

### D. Expansion to instance segmentation

mEMbrain’s output prediction until this step is a categorical image (*i*.*e*. each pixel is assigned to one of the classes the network was trained on) accompanied by its relative probability map (*i*.*e*. how sure the network is that a said pixel pertains to an assigned class). However, for the vast majority of connectomics tasks, each cell should be individually identifiable. Predictions of EM images in different classes are a powerful resource that can either strongly expedite manual reconstruction, or can be the first step necessary for many semi-automatic reconstruction methods. These labels can be directly imported in VAST and used in the following manners:

- **Machine learning-aided manual annotation with membrane-constrained painting** (i.e. “membrane detection+pen” mode). In this modality, the manual stroke of paint is restricted to be contiguous with mEMbrain’s membrane prediction. This allows the user to proceed in a swift manner, negligent of details such as complex borders that require a hefty amount of time if done precisely by hand.
- **Annotation with VAST’s flood filling functionality with underlying mEMbrain’s 2D segmentation**. By clicking once on the neurite of interest with the filling tool, the object is colored and expanded until it reaches the borders predicted by mEMbrain.
- Any other expansion algorithm that creates an instance segmentation starting from a semantic one.

## 4. Dataset Showcase

mEMbrain has been used to reconstruct neurons and neural circuits in a number of datasets, spanning different regions of the nervous system (including central and peripheral) at multiple scales (from cellular organelles to multi-nucleated cells) and across diverse species (including various invertebrates and mammals). Here we report some of the most interesting uses of mEMbrain insofar, showcasing a variety of unpublished datasets where our software had the opportunity to be tested, and where it played a pivotal role.

Importantly, we provide the ground truth datasets used to train networks for the segmentation cases here described. This should enable individuals who wish to train different architectures to bypass the time-expensive ground truth generation phase. Further, we also provide the trained networks that one can download for transfer learning purposes (showcased in Section F). The hope is that by providing pre-trained networks individual researchers will be provided with an advanced training starting point, and will be able to fine tune the network on their specific dataset with less ground truth needed, thus shortening the training times. Ground truth and networks are shared with the community at the following website: https://lichtman.rc.fas.harvard.edu/mEMbrain/

In all of the datasets, the predictions carried out by mEMbrain used a Nvidia RTX 2080Ti GPU, which computed at a speed of 0.2-0.25 seconds/MB. We assessed this range by recording the CPU time both prior to the point mEMbrain’s raw [1024×1024] pixels images are sent to the GPU (and before the MATLAB’s Deep Learning Toolbox memory-related operations) and again when the predictions are received on the mEMbrain’s endpoint (after the MATLAB’s Deep Learning Toolbox memory-related operations). To obtain the inference time on a dataset represented in VAST internally as [1024×1024] pixels tiles, one needs to linearly scale the above running time. We do not report on I/O and networking time because these widely vary on different architectures. Nonetheless, in the tests below the machine learning inference time significantly exceeded the I/O operations (using standard hard drive reading images at the order of 0.01 seconds/MB and writing at 0.05 seconds/MB). Indeed there are other factors related to running time that are not under mEMbrain’s management but are handled internally in VAST (as mentioned in Section C) and hence the running time in practice can be twice longer than the estimates reported here. For example, predicting the membranes on the whole mouse brain dataset (see Section A) took around 48 hours on a single desktop (see Section A for specifications) whereas inference time accounted for about 23.67 hours of the total running time. These running times adhere to the default network architecture used by mEMbrain (see Section B) and will vary accordingly for different architectures.

As a reminder, predictions can happen in different modalities with mEMbrain, such as by defining a box of the prediction with 3D start point and end point coordinates, or by defining a set of skeleton nodes in VAST (using VAST’s annotation layer) and letting mEMbrain follow these nodes for on-the-fly model predictions, or predicting on the whole dataset (see Section C). In all the showcases shown in this paper, one or many of these modes were used to assess the quality of the ground truth, allowing the researcher to quickly check the quality of the model performance on any sub-region of the large dataset or by applying sparse predictions around locations of interest. These methods also allowed easily revisiting regions predicted with earlier models when assessing the performance of a newly trained model. Whole volume predictions or predictions within a bounding 3D box were frequently used for the final prediction then used for reconstruction.

### A. The whole Mouse Brain dataset

We employed mEMbrain in our ongoing efforts to develop staining and cutting protocols that will eventually enable the reconstruction of a whole mouse brain (Lu *et al*., in preparation). In the current phase of the project, a newborn whole mouse brain was stained and cut, and several sections were stitched. The region of interest here shown is from the mouse’s motor cortex M2, covering layers II/III through VI. The sample was imaged with a Zeiss multibeam scanning electron microscope, at a resolution of 4×4×40 nm^3^/px, resulting in a total volume of 180×303×4 μm^3^.

The role of mEMbrain in this project was to assess the feasibility of reconstructing neural circuits when using such staining and cutting protocols. We started from a network pre-trained to detect cellular membranes on adult mouse cortex (32). We used this dataset for network pre-training because both datasets opted to preserve the extracellular space within the neuropil, using related staining techniques and resulting structure. Seven iterations of network training and manual corrections were needed in order to achieve good results, which amounted to 50 hours of ground truth preparation. The volume of the ground truth accounted for 5.8*/*10^4^ of the entire dataset. The criterion used to iteratively add ground truth and retrain the model was the appearance of faint membrane predictions or merge errors in the output of a 2-dimensional segmentation algorithm. The merge errors were detected by applying the network on all sections and a limited XY range using mEMbrain for the last iterations of network training. We used this approach only when it was already hard to detect possible membrane breaks in the output of the classifier. The added ground truth was selected by inspecting 2D segmentations from [2048 × 2048] pixels tiles from all sections in random locations. The majority of these errors appeared in the borders of cell bodies and occasionally due to tiny wrinkles incident to cell nuclei (a more detailed report of these dataset-specific considerations will appear in the relevant future publication). In total, ground truth included annotations from 339 distinct tiles and a total of 190MB of raw EM, of which 76% belonged to intracellular space and the rest to membrane and extracellular space. We then predicted all the cell membranes in the volume and segmented each 2D section. The predictions were carried out on a single desktop with a single GPU Nvidia RTX 2080 Ti, which required 5 days. Further, we used an automatic agglomeration algorithm (39) to reconstruct 3D cells; the high quality results with an exceptionally low rate of merge errors (see Figure 6), reassure that these new protocols may consent larger scale mouse brain reconstructions and will be discussed elsewhere (Lu *et al*., in preparation).

**Fig. 6.**
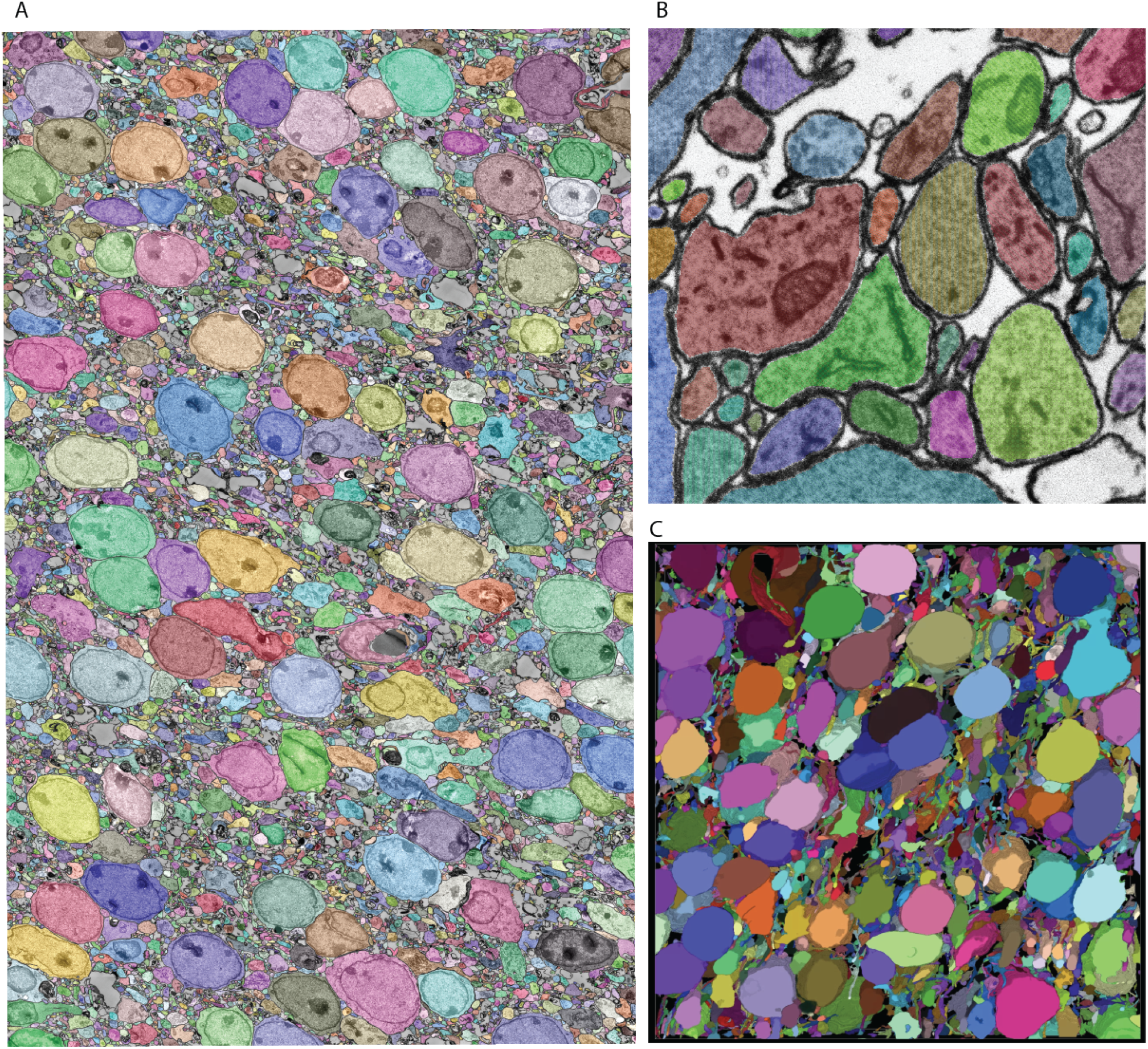
Showcase results of mEMbrain on the whole mouse brain dataset. **(A)**: 2D section of the whole mouse dataset segmented by using mEMbrain’s cell contour prediction in combination with automatic agglomeration methods (39). **(B)**: Example of a small region of interest of the dataset, meant to highlight the good quality of the results. **(C)**: Portion of a stack of sections, visualized in VAST’s 3D viewer. Lu *et al*., in preparation.

**Fig. 7.**
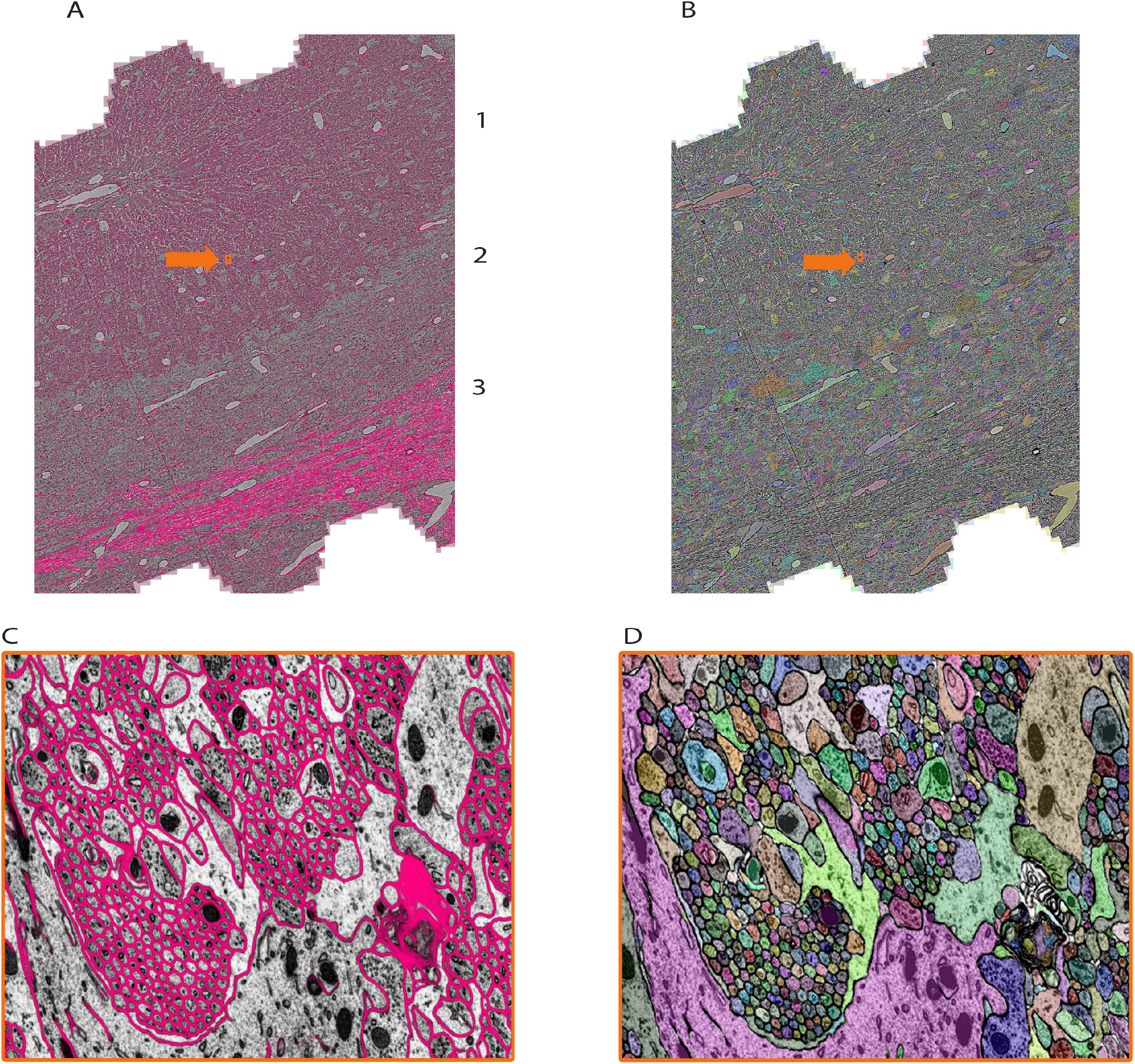
Showcase results of mEMbrain on the mouse cerebellum dataset from Dhanyasi et al., in preparation. **(A)**: predictions of membranes from mEMbrain overlaid on EM image. The layers of the cerebellar cortex are indicated: molecular layer (1), granule layer (2) and white matter (3). In orange, the location of the zoomed in panel in **C. B** and **D** showcase the same anatomy segmented instance-wise.

### B. The Mouse Cerebellum dataset

We tested mEMbrain on different regions of the mouse nervous system. Here, we report about our software’s use on the developmental mouse Cerebellum dataset (Dhanyasi *et al*., in preparation). The rationale behind this research is to study the development of the cerebellar circuits using electron microscopy. The region of interest is from the vermis, a midline region of the cerebellar cortex (60). The sample was imaged with a Zeiss multibeam scanning electron microscope at a resolution of 4×4×30 nm^3^/px, yielding a traceable volume of 650×320×240 μm^3^. To this end, mEMbrain processed and segmented 20% of this volume over a depth of 92 (3072 sections) and 20TB on disk.

In the context of this dataset, mEMbrain was used to expedite manual annotation, by both using mEMbrain’s predicted cell boundaries as constraints in VAST (see Section D, Method 1), and by carrying out 2D instance segmentation provided by our tool. As with the whole mouse brain dataset, also here we started the training from a network pre-trained to detect membranes on adult mouse cortex. Four iterations of network training and manual corrections were needed in order to achieve good results as inspected and assessed by the researcher on several key cell types and structures, which amounted to 60 hours of ground truth preparation. The volume of the ground truth accounted for 1.2/10^5^ of the entire segmented dataset. Iterations proceeded as long as merge errors were manually detected in randomly selected tiles across the entire dataset. Special attention was given to the predicted membranes in Purkinje cell dendrites and the parallel fibers innervating them, while prediction quality in the white matter and the granule cell bodies was not assessed (as these escaped the research goals). In total, ground truth included annotations from 33 tiles and a total of 286MB of raw EM, of which 78% belonged to intracellular space and the rest to membrane and extracellular space.

The researcher reported that the greatest speed-up for this dataset was provided by the 2D instance segmentation. To corroborate this assessment, an additional speed test was performed by three annotators, and an estimate of the expedition offered by several semi-automatic methods is recounted in Section 5.

### C. The *C. elegans* dataset

We assessed our software on a number of invertebrates. Here we show mEMbrain’s employment on one *C. elegans* dataset. This sample (58) was a wildtype nematode in the dauer diapause, an alternative, stress-resistant larval stage geared towards survival (61). The sample, with a cylindrical shape in a diameter of 15.8 μm was imaged with a focused ion beam - scanning electron microscope (FIB-SEM) at a resolution of 5×5×8 nm^3^/px (58). A region including 2998 serial sections (24 μm), containing the nerve ring, surrounding tissue and the specimen’s body wall, was cropped for connectomic segmentation.

For this dataset, mEMbrain was used as a semi-automatic segmentation tool. We started the training from a network pre-trained to detect membranes on the octopus vertical lobe described below in more detail. Five iterations of network training and manual corrections were needed in order to achieve satisfying results as evaluated on the basis of the membrane appearance in the nerve ring and the surrounding neuron and muscle cell bodies. The ability of the predicted membranes to separate cells was manually tested on suspected regions by flood-filling the membrane probabilities in 2D using VAST’s functionality on selected regions (48). The researchers computed that semi-automatic ground truth generation cut the manual annotation labor time by a little less than 50%: the ground truth required for the first training iteration took 14 hours of manual annotation. Similarly, also subsequent iterations cumulatively required 18 hours of painting. However, the volume traced in this amount of time is doubled with respect to the first iteration. The workflow of this dataset is shown in Figure 2. In total, ground truth included annotations from 42 tiles and a total of 37MB of raw EM, of which 70% represented intracellular space of neurons, glia and muscles and the rest accounted for other tissues, the body wall as well as a representation of the imaged regions exterior to the worm. The latter was needed to avoid erroneous merging of cellular objects with the exterior, leading to large merge errors among neurons close to the cell body of the animal.

### D. The Octopus Vertical Lobe dataset

We had the unique opportunity to test mEMbrain on non-conventional model organisms in the neuroscience community, thus testing the usefulness and generalizability of our tool across species. In particular, we were excited to assess mEMbrain on a sample from the *Octopus vulgaris* dataset (18). The region of interest is in a lateral lobule of the *Octopus vulgaris*’ vertical lobe (VL), a brain structure mediating acquisition of long-term memory in this behaviorally advanced mollusc (62, 63). The sample was imaged at high resolution with a Zeiss FEI Magellan scanning electron microscope equipped with a custom image acquisition software (64). The ROI was scanned over 891 sections each 30 nm thick at a resolution of 4 nm/px, constituting a traceable 3D stack of 260×390×27 μm^3^.

mEMbrain was here mostly used for aiding manual annotation. We started the training from scratch without pre-training be-cause this dataset is the first volumetric analysis of ultrastructure of the octopus central brain (18). Four iterations of network training were needed in order to achieve satisfying quality which was evaluated based on the appearance of predicted membranes in two different regions of the neuropile (contacts between the input axons to the Amacrine interneurons and contacts between Amacrine neurons and Large neurons; (18)). The first ground truth annotation already provided good network for most of the neuropile and the two other iterations were needed in order to improve the quality of glial processes and cell bodies. The dataset included broken membranes for the main trunk of the large neuron for unknown reason. Predicted membranes were not satisfactory for 2D and 3D for these processes. In total, ground truth included annotations from 176 tiles and a total of 761MB of raw EM (1.5/10^4^ of the entire segmented dataset).

As described in Section D, the output semantic segmentation obtained with mEMbrain can be directly utilized in VAST as constraints for the annotation of objects. In this manner, a single drop of paint floods the entirety of the neurite, and allows the researcher to proceed in a swift manner, without needing to pay attention to anatomical details. For this dataset, the researchers using our software reported that there is a 2-fold increase in speed with mEMbrain’s aid when the purpose is to simply roughly skeletonize a cell, not being mindful of morphological details. However, the most significant advantage of using mEMbrain is the expediency of precise anatomical reconstructions, given that accurate reconstructions consume a sizeable amount of manual time. Instead, with mEMbrain, the time to skeletonize a neurite matches the time it takes to reconstruct it accurately; explaining why in this modality there is a 10-fold increase in speed when using mEMbrain. For example, this allowed for a fast and precise reconstruction of axonal boutons and cell bodies, which enabled subsequent morphometric analysis (see Figure 8).

**Fig. 8.**
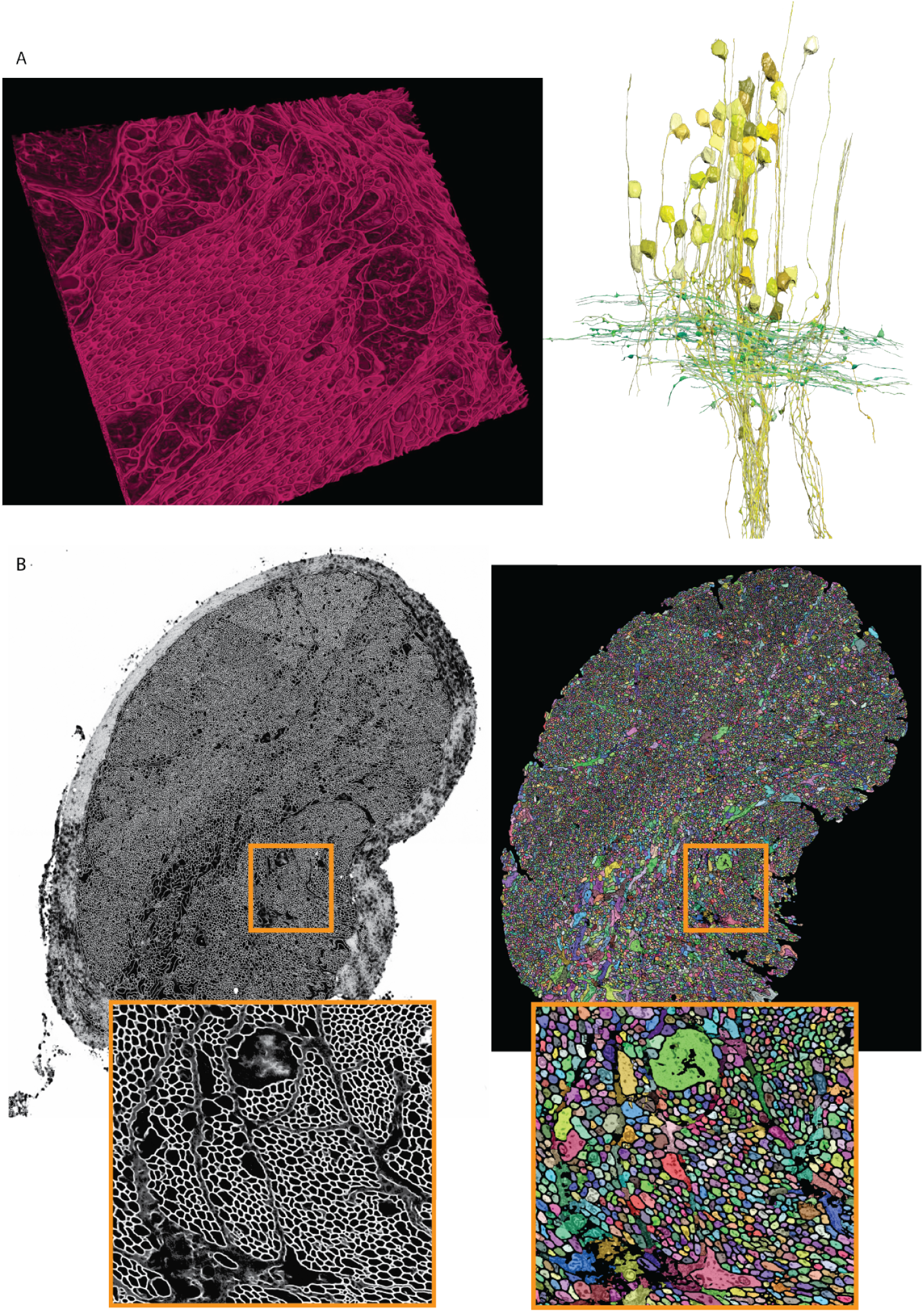
Showcase results of mEMbrain on the molluscs’ datasets. **(A):** Examples of the *Octopus vulgaris* dataset, from (18). On the left, sample of the cell boundaries predicted by mEMbrain and shown in VAST’s 3D viewer. On the right: 3D rendering of interneurons (yellow) and afferents (green) in the learning and memory brain center in the octopus brain. Reconstruction mode: pen annotation constrained “on-the-fly” in VAST by mEMbrain’s border probabilities. **(B):** Examples of the dataset from the rhinophore connective of the nudibranch, *Berghia stephanieae* (59). The membrane predictions (left) and the instance segmentation (right) are shown for a whole connective slice; the instance segmentation was obtained starting from mEMbrain’s cell boundaries and applying a 3D agglomeration algorithm (39). To appreciate the sheer number of processes connecting the brain to the rhinophore, small regions are zoomed out in orange.

### E. The *Berghia stephanieae* dataset

We tested our tool on a second mollusc, the nudibranch *Berghia stephanieae*, a species of sea slug newly introduced for neuroscience research. The aim of this project is to determine the synaptic connectivity of neurons in the rhinophore ganglion, which receives input from the olfactory sensory organs. The rhinophore connective contains axons that travel between the rhinophore ganglion and the cerebral ganglion. The sample of the rhinophore connective here was sectioned at 33 nm and imaged with a Zeiss scanning electron microscope at a resolution of 4 nm/px (59), yielding a traceable volume of 134×41×1 μm^3^.

The *Berghia* dataset was the first one on which we witnessed the power of transfer learning (see Section F). Seven-ten hours of ground truth annotation produced a handful of labels from 31 tiles and a total of 11.2 MB of raw EM images, that were used to perform continuous learning from a network pre-trained on the *Octopus vulgaris* dataset. This dataset demonstrates the usability of pre-trained networks in cases where the target dataset has a very limited amount of ground truth. mEMbrain’s output was used to obtain 3D segmentation when agglomerated with automatic algorithms (39). This reconstruction enabled the possibility to automatically count the number of processes present in the rhinophore connective tissue region, and revealed that this part of the nudibranch nervous system harbors an exceedingly high number of processes (roughly 30 000 - the counting was double checked by manual inspection). This was an important finding, as the *Berghia stephanieae*’s rhinophore ganglion itself contains only 9000 cell bodies (59). The complex organization and the abundance of processes (shown in Figure 8) suggest that such peripheral organs are highly interconnected with the central nervous system of the animal, sharing similarities with octopuses and other cephalopods (65, 66).

### F. Transfer Learning

One tool that we found incredibly valuable in our reconstructions was using knowledge learnt from one dataset and applying it towards others, leveraging the concept of *transfer learning*, and more specifically of domain adaptation (67). We experimented with a variety of modalities for transfer learning. We started by freezing all the model’s weights except for the last layer, a strategy that maintains the internal representations previously learned by the model, while fine tuning the last layer for the specific new dataset at hand. We then tested the idea of freezing only the model’s encoding weights, in other words the first half of a U-Net architecture, while allowing the decoder’s weights to fine tune for the new dataset. Further, we explored allowing the encoder to learn at a very slow rate (maintaining most of the pre-trained knowledge), typically 10 times smaller than the decoder’s learning rate, in a technique called “leaky freeze”. Moreover, we tested applying a continuous learning approach, whereby after training on a first dataset, the same network is trained on a second one without modification of its learning rates. One concern that might arise with this approach is the occurrence of catastrophic forgetting, which is the tendency of a network to completely and abruptly forget previous learned information, upon learning new information (68). For this reason, we also tested an episodic memory strategy, where the training schedule interleaves learning from the two datasets at hand.

The main conclusion of our multiple experiments is that the strategy of transfer learning significantly reduces the time needed to achieve satisfactory results; pre-trained networks have already learned multiple fundamental features of EM images, tentatively distinguishing membranes of cells. Thus, the training of networks on subsequent datasets is geared towards fine-tuning their *a priori* knowledge and adapting it to the specific dataset at hand. This means that the number of epochs - that is the number of passes of the whole training dataset that the deep learning network has completed - required for good performance is significantly less than when training a network from scratch. Furthermore, the amount of ground truth needed to achieve satisfactory results is also drastically reduced, as many of the features - such as edge detection, boundary detection, and general interpretation of different gray scales of electron microscopy images - have already been assimilated from learning on the previous data. The second conclusion from our tests highlights that the strategy of continuous learning is the one that yielded the best results. Further, this method is particularly user-friendly given that no alterations to the network need to be made.

It is important to note that transfer learning works best when the network trains on datasets that share many common features. One striking example where transfer learning proved to be a powerful technique was in the *Berghia stephanieae* dataset. For this project, the human-generated ground truth was reasonably scarce, and hence when a network was trained with mEMbrain for semantic segmentation, the outcomes were quite poor, as can be seen in Figure 9. However, we noticed a qualitatively strong resemblance between the EM image properties of the *Berghia stephanieae* and of the *Octopus vulgaris*. We reasoned that this could be a case in which transfer learning techniques would be especially impactful in aiding the paucity of ground truth to learn from. Thus, we took the best-performing network trained on the *Octopus vulgaris* and we trained it in a continuous learning fashion for 5 subsequent epochs on 3 ground truth images from the *Berghia stephanieae* dataset. Within only 10 minutes of training, the validation accuracy of the network reached 97% and the results were of high quality, as can be seen from Figure 9. Hence, working with pre-trained networks and fine-tuning them on the specific dataset at hand dramatically reduces the time invested both in ground truth generation and in training of the network. We highly recommend to save previously trained networks and to further their learning on new datasets in order to expedite the segmentation process.

**Fig. 9.**
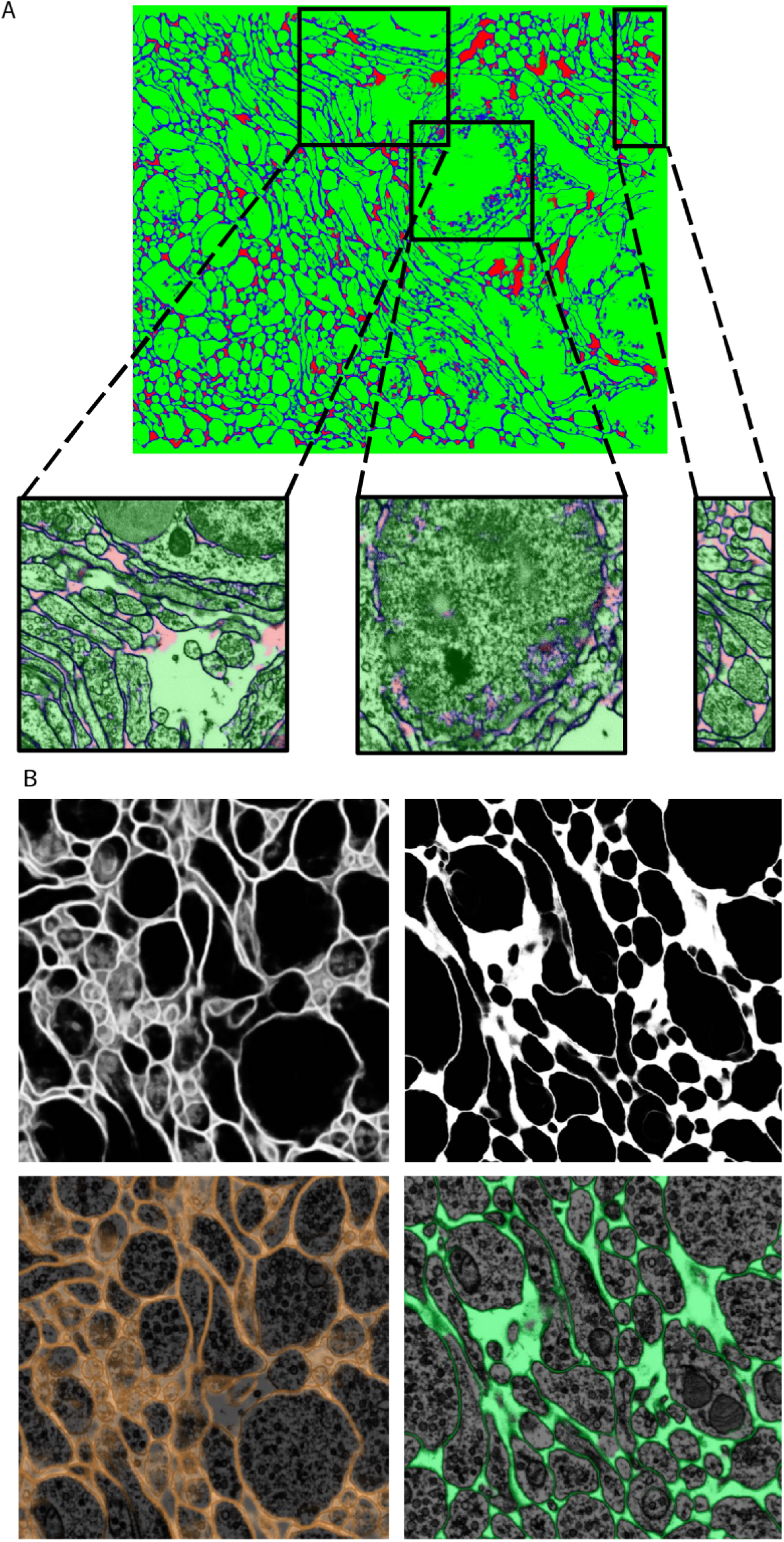
Transfer learning approaches from the Octopus to the Berghia stephanieae datasets. **(A)**: example of poor generalization of the network on the *Berghia* dataset, due to limited training ground truth. In green the intracellular space, in blue the cell boundaries, and in red the remainders. **(B):** networks pretrained on the *Octopus* dataset predictions on the *Berghia* dataset without continuous learning (left) and with continuous learning (right). In grayscale are membranes only, while below they are overlayed to EM images.

## 5. Evaluating Speed Up with Machine Learning-Aided Painting

We tested the speed up provided by mEMbrain’s output by conducting a proof-of-concept timed experiment. We asked three experienced researchers to manually annotate one neurite for 10 minutes. We then compared the resulting labeled volume with the volumes annotated by the same researchers when using mEMbrain’s output in combination with VAST’s tools. In particular, we tested:

- using mEMbrain’s 2D segmentation in combination with VAST’s pen annotation mode (Section D, Method 1);
- using mEMbrain’s 2D segmentation in combination with VAST’s filling tool (Section D, Method 2);
- using machine learning-aided manual annotation with membrane-constrained painting carried out with VAST’s pen annotation mode;

We benchmarked such methods against manual annotations only. The tests were carried on the *Mouse Cerebellum* dataset, presented in Section B. The results are quantified in Figure 10C. The main finding is that painting with an underlying machine learning aid is at least 20 times faster than labeling purely with manual approaches. More specifically, the combination of mEMbrain’s 2D segmentation together with VAST’s pen annotation model yields the fastest results, particularly when striving for accuracy. In contrast, opting for mEMbrain’s 2D segmentation in tandem with VAST’s flooding tool, while vastly accelerating manual labor, might be suboptimal in scenarios in which VAST’s flooding tool could yield to merge errors, which in turn require more time for correction and label postprocessing. However, this modality has been reported by our user to be most ergonomic. This speed evaluation will need to be corroborated by future tests on different datasets.

**Fig. 10.**
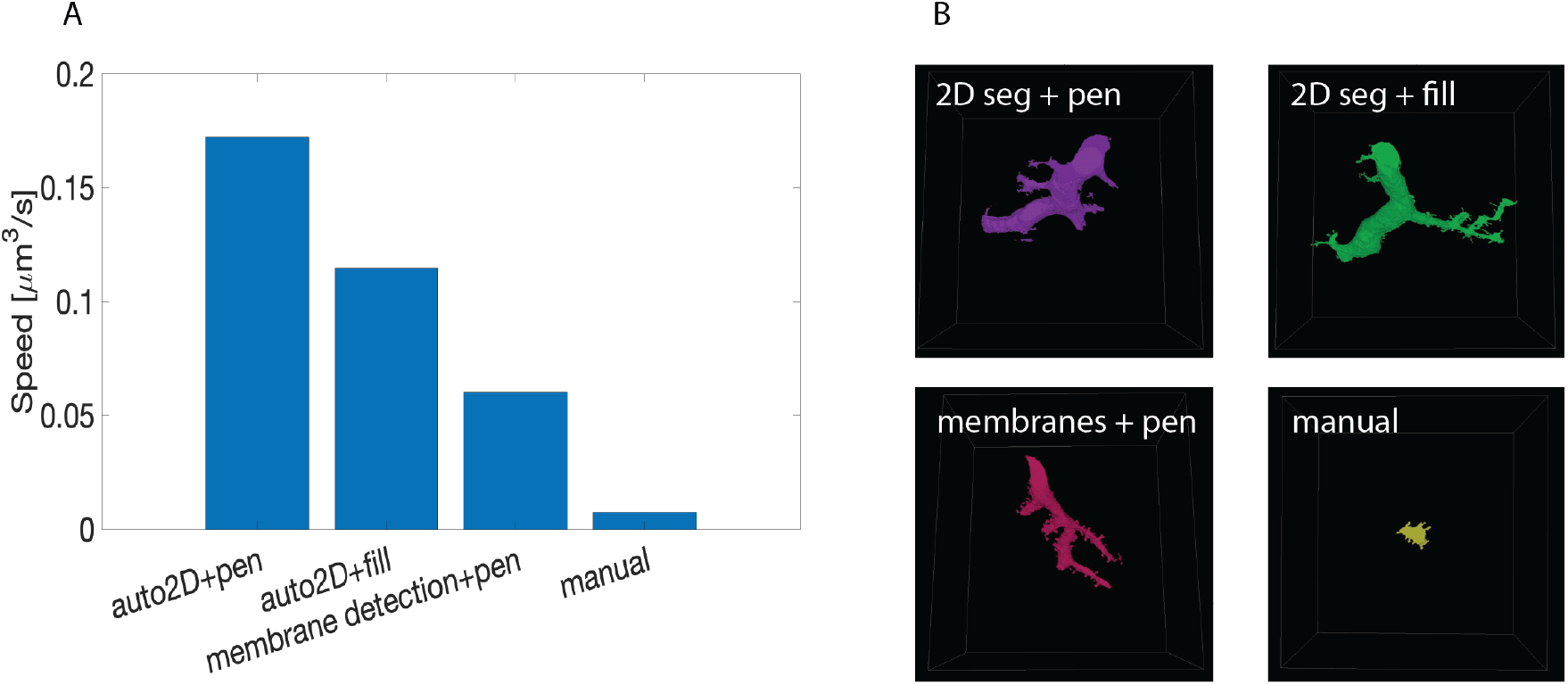
Summary of speed up test with machine learning-aided painting. **(A)**: bar graph of the average speed of three machine learning-aided painting modalities together with manual annotation. The test was performed by 3 experienced annotators on the Cerebellum dataset. **(B):** sample visual results of the speed test from one annotator painted in 10 minutes. Top left: volume painted with the 2D segmentation in tandem with VAST’s pen mode. Top right: volume painted with the 2D segmentation together with VAST’s fill mode. Bottom left: volume painted when membrane detections are used with VAST’s paint mode. Bottom right: volume painted with manual annotation only.

## 6. Comparison with other tools

While there are many free software tools in the field for labeling and manual annotation, visualization and proofreading, there are fewer software providing a comprehensive and user-friendly pipeline for CNN training geared towards EM segmentation. One first aspect to notice is that all software, mEMbrain included, rely on other packages for visualization and proofreading. The power of mEMbrain relies precisely in its synergy with VAST, which is excellent for data handling, visualization, annotation, and offers a variety of tools that can be co-leveraged together with our software. For these reasons, mEMbrain features the very useful ability to predict on-the-fly in regions chosen by the researcher and immediately visualizable in VAST. This greatly enables the scientist to assess the quality of mEMbrain’s outcome, and mitigates the the time for import and export of datasets and segmentations.

Another feature of mEMbrain we deem fundamental is its wrapping of all the pipeline in one unique GUI, without the user having to interact with code and having to master different interfaces. Importantly, mEMbrain provides the ability to create datasets necessary for the training phase, which are data-augmented in order to enhance the learning abilities of the network. In Table 2 we show a brief summary of the salient points we reckoned important for a user-friendly software tool compared across the packages most similar to mEMbrain.

**Table 1.** Information about the ground truth here released to the community.

| Dataset | Amount of ground truth | Hours invested in generating ground truth | Paper to look out for more information |
| --- | --- | --- | --- |
| *Whole mouse brain* | 339 tiles, 190 MB | 50 | Lu et al., in preparation |
| *Cerebellum* | 33 tiles, 286 MB | 60 | Dhanyasi et al., in preparation |
| *C. elegans* | 42 tiles, 37 MB | 32 | (58) |
| *Octopus vertical lobe* | 176 tiles, 761 MB | 50-60 | (18) |
| *Berghia stephanieae* | 31 tiles, 11.2 MB | 7-10 | (59) |

**Table 2.**
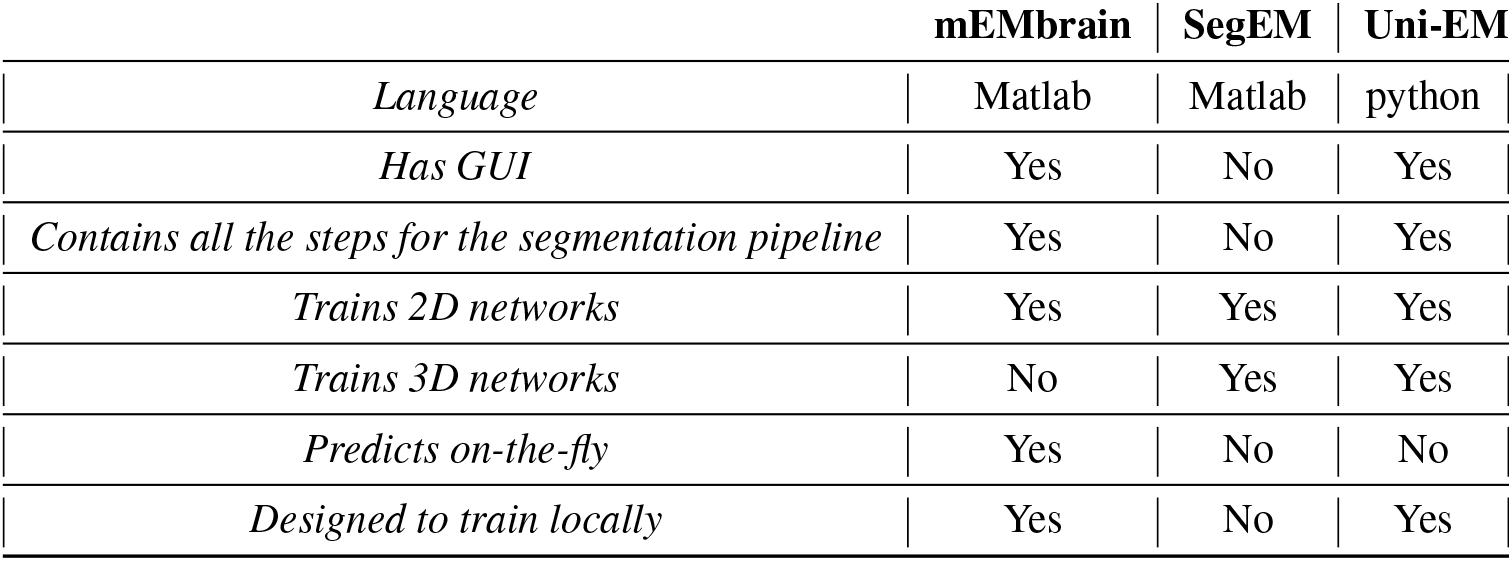
Summary of the comparison between the state-of-the-art (semi) automatic segmentation pipelines in connectomics. The qualities in the various rows represent some of the parameters we deemed important when designing mEMbrain.

|  | mEMbrain | SegEM | Uni-EM |
| --- | --- | --- | --- |
| *Language* | Matlab | Matlab | python |
| *Has GUI* | Yes | No | Yes |
| *Contains all the steps for the segmentation pipeline* | Yes | No | Yes |
| *Trains 2D networks* | Yes | Yes | Yes |
| *Trains 3D networks* | No | Yes | Yes |
| *Predicts on-the-fly* | Yes | No | No |
| *Designed to train locally* | Yes | No | Yes |

## 7. Discussion and Outlook

Here, we presented a software tool - mEMbrain - which provides a solution for carrying out semi-automatic CNN-based segmentation of electron microscopy datasets. Importantly, our package installation is straightforward and limited to the download of a folder, and it assumes little to no prior coding experience from the user. mEMbrain works synergistically with VAST, a widely used annotation and segmentation tool in the connectomics community (48). Our hope is that VAST users will be enabled in their reconstructions thanks to mEMbrain.

Our tool compares favorably to other similar published software tools. One feature that we hope to incorporate in future editions of mEMbrain is the possibility to train on state-of-the-art 3D CNNs, such as 3C (39), thereby allowing for better results. Nevertheless, it is important to note that 2D section segmentation can provide satisfactory results, depending on the quality of the sample staining and the dataset alignment.

One of the main motivations for coding mEMbrain was its capability for processing datasets and running deep learning algorithms on local computers. Although at first sight this may appear as a set-back, it represents a tangible means for affordable connectomics by abolishing the costs for expensive clusters. Furthermore, it avoids the need of transferring massive datasets in different locations, which results in a gain in terms of time, and allows for a rapid validation of results due to its close dialogue with VAST. Many of the results showed in this paper were obtained by using a single Nvidia GPU RTX 2080 Ti. Thus, with the current technology the use of mEMbrain is best when the dataset is within the terabyte range. To this end, a useful extension of the toolbox would be to allow the possibility of predicting on a computing cluster when the user necessitates it. Moreover, we noticed how the main bottleneck of the predicting time is created by MATLAB reading chunks of data from VAST. Therefore another possible future direction is to allow for the prediction of multiple classifiers at the same time (e.g. co-prediction of mitochondria and vesicles) in order to avoid reading the dataset multiple times. Nevertheless, it is foreseeable that the available technology will improve, and with it also the prediction time with mEMbrain.

Finally, from the locality of our solution stems the exciting opportunity to place the segmentation step of the connectomics pipeline next to the scope, and to readily predict each tile scanned by the electron microscope, allowing researchers to access their on-the-fly reconstruction in a more timely fashion (5).

## Author Contributions

E.C.P., Y.M. and J.W.L. conceptualized the idea of mEMbrain. E.C.P. and Y.M. coded the first prototype of the software. E.Y. and Y.M. wrote the code for the current version of the software. F.B., B.H., X.L, N.D., Y.M., M.W., M.Z., B.D., P.S.K. and F.Y. generated ground truth, tested and evaluated mEMbrain on different datasets, provided figures for the manuscript. N.D. conducted the machine learning speed up test and Y.M. analyzed the results. C.F.P. ran the tests benchmarking mEMbrain’s patch generation with data augmentation method against off-the-shelf methods under supervision of A.D.T.S.. M.B. pioneered the work on the skeleton network under supervision of Y.M.. E.C.P. wrote the manuscript, with input and help from all the other co-authors.

## Data Availability Statement

The code generated for this study can be found in the mEMbrain repository [https://github.com/emmay78/mEMbrain].

## ACKNOWLEDGEMENTS

The authors would like to thank Anna Steyer, Sebastian Britz, Sebastian Markert, Yannick Schwab and Christian Stigloher for generating the *C. elegans* FIB-SEM dataset. The authors thank Daniel R. Berger for his assistance in interfacing MATLAB codes with VAST.

