## Supplementary Material for "mEMbrain: an interactive deep learning MATLAB tool for connectomic segmentation on commodity desktops"

### 1. Supplementary Data

#### A. Installation

One of the advantages of mEMbrain is it's nearly instantaneous usability, given that its installation is straightforward and limited to a folder download. We ask the users to first make sure they have MATLAB installed on the computer they intend to use mEMbrain on. In particular, the users should make sure to have installed with MATLAB also the Computer Vision Toolbox, the Deep Learning Toolbox and the Parallel Computing Toolbox, and agree to install whatever dependencies these may have. If done from scratch, MATLAB's installation could take up to 30 minutes (often less).
After verifying that MATLAB is running, one can download mEMbrain's folder of code from https://github.com/emmay78/mEMbrain. The folder consists of 352 KB, and thus downloads quasi instantaneously. Once downloaded, mEMbrain's folder should be moved in an appropriate location on the computer. To get started, it is sufficient to double click on the $mEMbrain_GUI.mlapp$ file; this will launch one of mEMbrain's main user interfaces which collects dataset generation and preprocessing, network training and prediction from VAST.
To maximize mEMbrain's potential, it is advisable to work with the software VAST for annotation, segmentation and visualization. Currently, VAST is available to Windows users. To download VAST, proceed to https://lichtman.rc.fas.harvard.edu/vast/ VAST, as mEMbrain, also has a straightforward: it comes as zipped folder of code. To launch, simply unzip the folder and launch the executable (3.4 MB). This websites also provides some sample datasets with which the user might want to test mEMbrain out.

### B. mEMbrain as part of 3D instance segmentation pipelines

Although currently mEMbrain does not have a 3D pipeline incorporated within its GUI, nevertheless our tool has been used for enabling 3D reconstructions. In fact, many of the pipelines for 3D reconstruction do rely on having high quality membrane prediction, which is one of our software's main output. Thus, we tested mEMbrain as an essential prerequisite for one of the published 3D pipelines, namely the Cross Classification Clustering (3C) algorithm (34), and its results are then showcased in the *C. elegans* and the whole mouse brain datasets in Section 4.

With a few exceptions, all of the widely used reconstruction pipelines make an intermediate use of a membrane probabilities and an additional provisional over-segmentation (Turaga et al., 2010; Lee et al., 2015; Meirovitch et al., 2016; Lee et al., 2017; Wolf et al., 2018; Beier et al., 2017; Funke et al., 2018; Meirovitch et al., 2019; Macrina et al., 2021).

We demonstrate the utility of this approach by leveraging recent agglomeration techniques, such as Cross Classification Clustering (34), combined with novel predictions of neuronal backbones, described in Section 3.1. In a first step, mEMbrain's output inferences are used to partition the space. This first over-segmentation, which aims at minimizing the number of objects internally contained in a neuronal boundary, is subsequently merged. While some pipelines attempt to achieve this goal with small 3D supervoxels (Macrina et al., 2021), we used here 2D segmentations that have a good representation of the neuronal cross section (Figure 4.2 A). Agglomeration proceeds with an optimization step that matches 2-D objects across sections. This process may be aided and furthered by a novel agglomeration technique based on supervised learning of the medial axis of neurons, or 2-dimensional skeletons, as we presented in Section 3.1, which are learned from the conversion of membrane ground truth to skeleton ground truth. Our tests on the various datasets (see Section 4) demonstrate such convolutional neural networks can learn the medial axis of thin neurites and the intricate morphology of enwrapping glial cells, leading to complex topology of the skeletons such as circular skeletons for enwrapping objects and confident detection of bifurcations, including spine protrusions from dendritic shafts (see Figure 1). Overall, our empirical tests propose that learning and predicting skeletons directly from the EM images provides additional information to traditional pipelines. In particular, the agglomeration step can make use of such information by assembling objects whose skeleton strongly overlap across planes. Results of this algorithm are visible in Figure 6.

### 2. Supplementary Tables and Figures

#### A. Figures

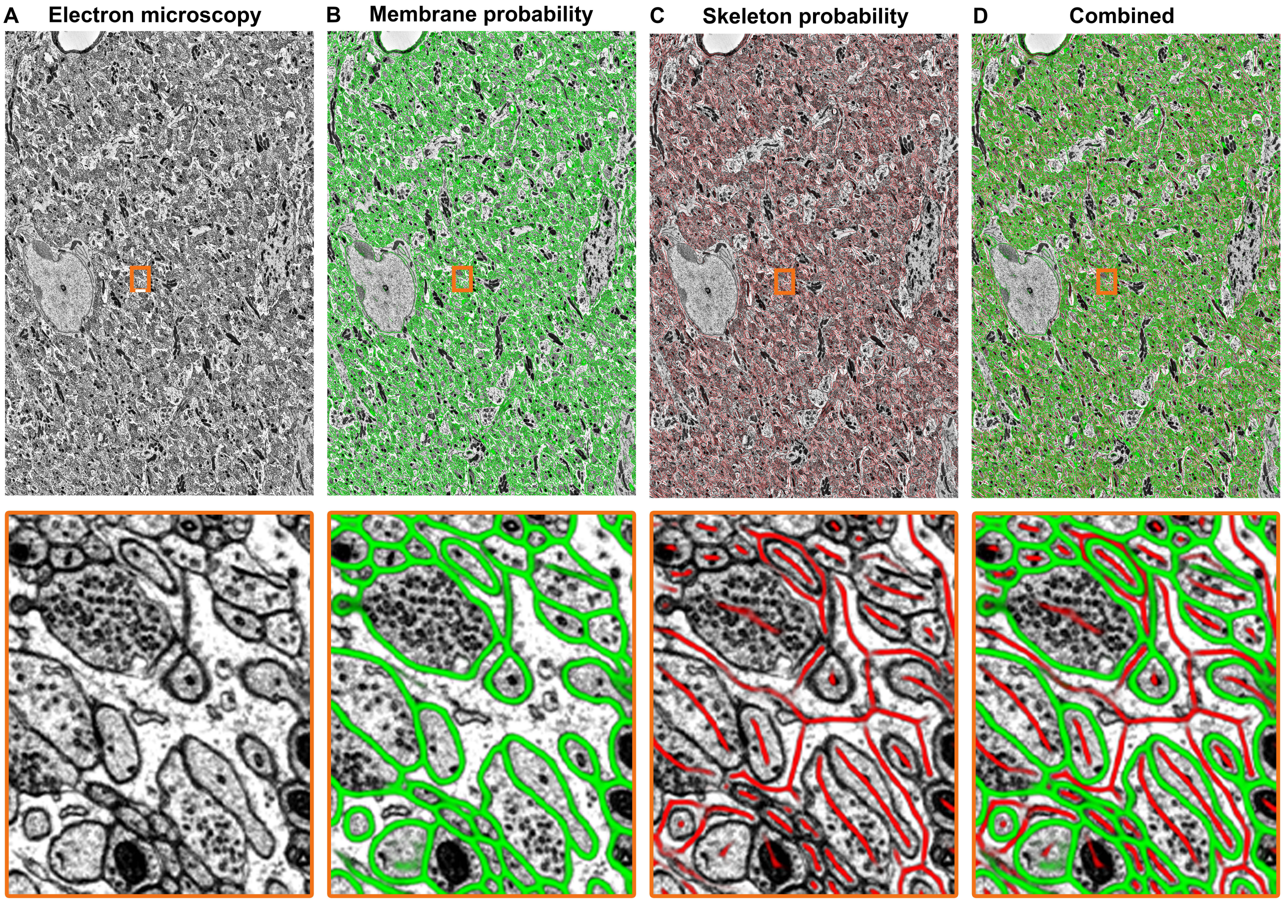


**Fig. 1.** *mEMbrain's skeleton predictions.* **(A)**: Electron microscopy tile from from the Cerebellum dataset (see Section 4.2). Dimensions: 5731 x 3813 pixels. Resolution: 8nm per pixel along the x and y axes. **(B)**: mEMbrain's membrane predictions achieved with supervised learning using expert-made membrane ground truth. **(C)**: Skeleton predictions achieved with supervised learning. The ground truth of such learning is provided by mEMbrain's automatic conversion of expert-made membrane ground truth into skeleton ground truth (see Section 3.1). Similar to the membrane neural network, the input to the skeletonizing network is the EM image with 4nm pixel resolution. **(D)**: Superposition of the membrane and skeleton probabilities as provided by the two separately trained convolutional neural networks. The magnified images show the ability of the skeleton predictions to correctly detect continuous structures - often glia - that enwrap around neurons, and thin neurites whose membrane probabilities are not always neatly defined.

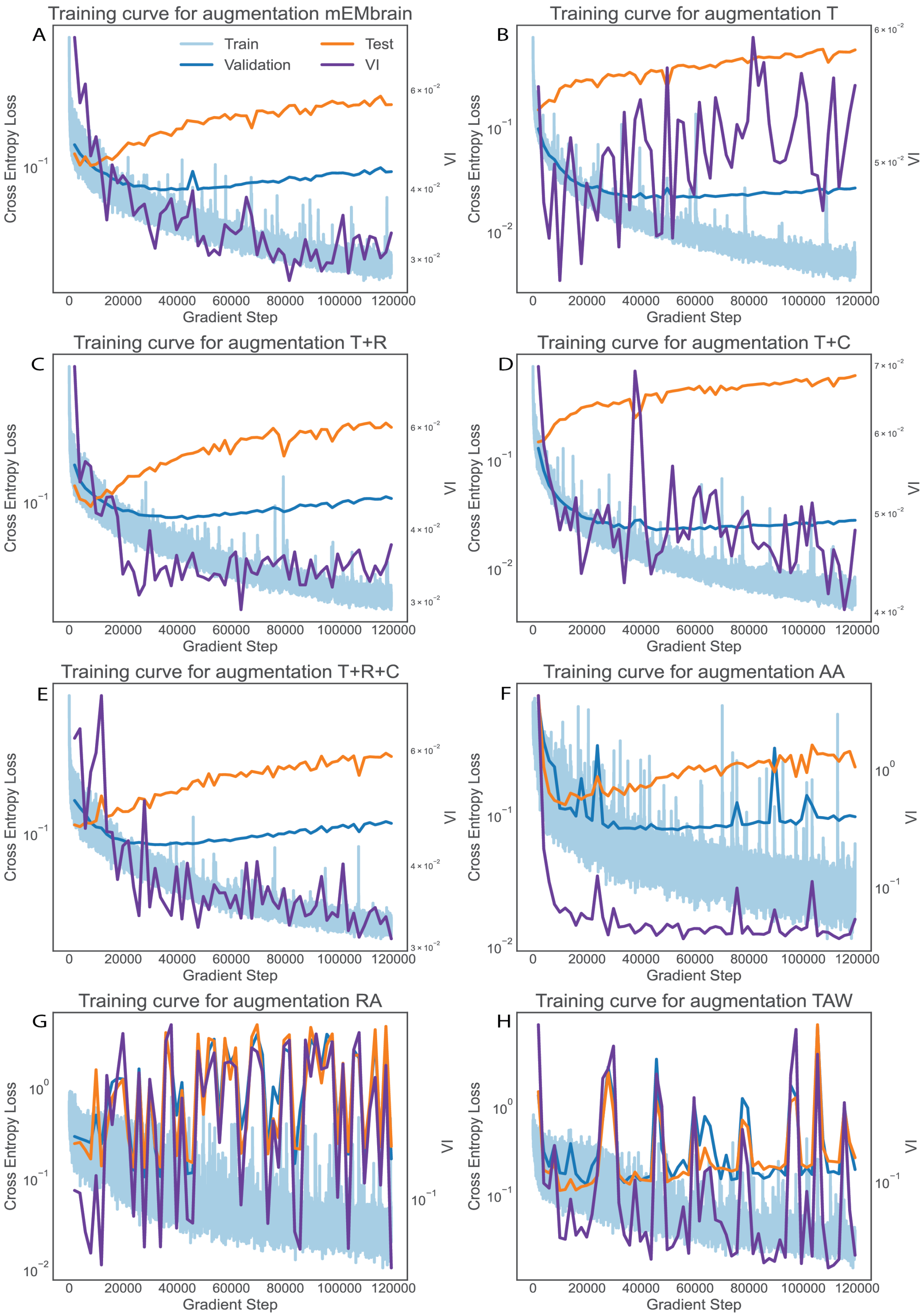


**Fig. 2.** Training loss, validation loss, test loss and test variation of information (total VI) for different augmentation methods. The augmentation methods are as follow. (A): mEMbrain. (B): translation only. (C): translation and rotation. (D): translation and color jitter augmentation. (E): translation, rotation and color jitter augmentation. (F): AutoAugment (Cubuk et al., 2018). (G): RandAugment (Cubuk et al., 2020). (H): Trivial Augment Wide (Muller et al., 2021). We observe that mEMbrain achieves a lower test total VI compared to simple baselines and published general methods. It is also worth noting that the test total VI decreases even after the validation and test loss ceases to improve.

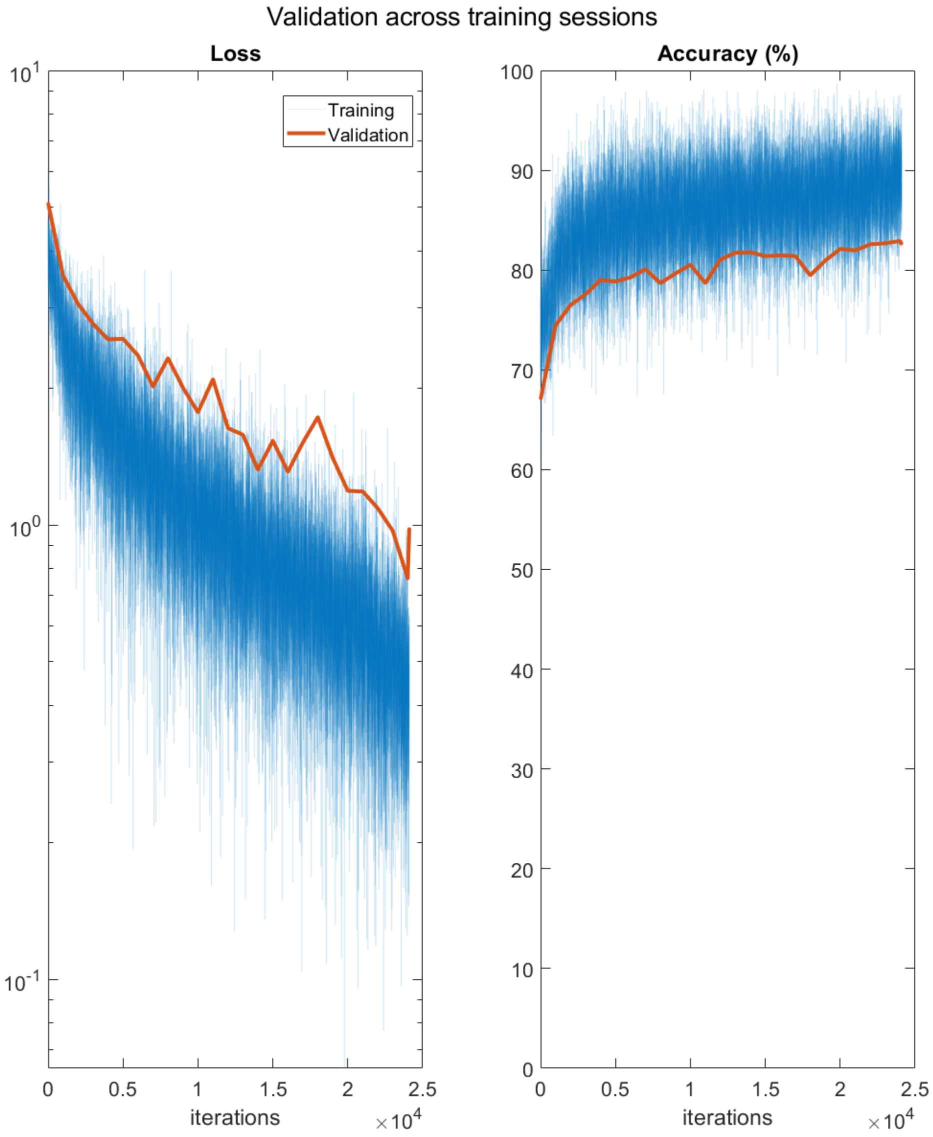


**Fig. 3.** *Example training and validation curves.* The ground truth used was from the Cerebellum dataset, described in Section 4.2. The validation data was obtained posterior from an independent ground truth, using EM images that are wrongly segmented by previous iterations and by the network being trained. The validation curve continues to drop which signifies that in this training regime over-fitting does not occur, which implies that the network's accuracy is not saturated. In agreement with the qualitative assessment (see figure), the gap between the validation accuracy and the training accuracy indicates that adding training iterations with the validation ground truth will increase the expressibility of the model.

### B. Tables

| Hyperparameter | Default value and brief explanation |
|---|---|
| U-Net Depth | Default value: 5<br>It determines how many times the input image will be downsampled/upsampled in the U-Net's encoding/decoding phase.<br>For more information about this parameter, please<br>visit https://www.mathworks.com/help/vision/ref/unetlayers.html#mw_6f8d62d2-0e3a-4188-aee0-0919868c41d3 |
| Initial Learning Rate | Default value: 0.00005<br>Roughly speaking, it determines how fast the network will learn.<br>Setting a very high learning rate may run in the risk of overshooting the optimal solution,<br>while setting a low one may yield a slow learning process. |
| Training Optimizer | Default: Adam<br>Other possibilities are: Stochastic Gradient Descent (SGD), and Root Mean Square Propagation (RMSProp).<br>All three are iterative algorithms used to optimize an objective function. |
| Training/Validation Split | Default: 0.7<br>This proportion describes how much data the user chooses to allocate towards training vs validation. |

**Table 1.** Table providing a summary of the key hyperparameters involved in (U-Net) network training, together with a simplistic explanation of their meaning and the default value in mEMbrain. Importantly, the default parameters are not necessarily a one-size-fits-all optimal solution. These are values that we used in our own research and have generally worked well. The user should feel free to experiment and change the parameters at will to explore the best values for their own research project.
